# Mitochondrial protein C15ORF48 is a stress-independent inducer of autophagy that regulates oxidative stress and autoimmunity

**DOI:** 10.1101/2023.10.17.562644

**Authors:** Yuki Takakura, Moeka Machida, Natsumi Terada, Yuka Katsumi, Seika Kawamura, Kenta Horie, Maki Miyauchi, Tatsuya Ishikawa, Nobuko Akiyama, Takao Seki, Takahisa Miyao, Mio Hayama, Rin Endo, Hiroto Ishii, Yuya Maruyama, Naho Hagiwara, Tetsuya J. Kobayashi, Naoto Yamaguchi, Hiroyuki Takano, Taishin Akiyama, Noritaka Yamaguchi

**Affiliations:** Department of Molecular Cardiovascular Pharmacology, Graduate School of Pharmaceutical Sciences, Chiba University, Chiba 260-8675, Japan; Laboratory of Molecular Cell Biology, Graduate School of Pharmaceutical Sciences, Chiba University, Chiba 260-8675, Japan; Laboratory for Immune Homeostasis, RIKEN Center for Integrative Medical Sciences, Yokohama 230-0045, Japan; Immunobiology, Graduate School of Medical Life Science, Yokohama City University, Yokohama 230-0045, Japan; Institute of Industrial Science, The University of Tokyo, Tokyo 153-8505, Japan

## Abstract

Autophagy is primarily activated by cellular stress, such as starvation or mitochondrial damage. However, stress-independent autophagy is activated by unknown mechanisms in several cell types, such as thymic epithelial cells (TECs). Here we report that the mitochondrial protein, C15ORF48, is a critical inducer of stress-independent autophagy. Mechanistically, C15ORF48 reduces the mitochondrial membrane potential and lowers intracellular ATP levels, thereby activating AMP-activated protein kinase and its downstream Unc-51-like kinase 1. Interestingly, C15ORF48 induction of autophagy upregulates intracellular glutathione levels, promoting cell survival by reducing oxidative stress. Mice deficient in *C15orf48* showed a reduction in stress-independent autophagy in TECs, but not in typical starvation-induced autophagy in skeletal muscles. Moreover, *C15orf48*^−/–^ mice developed autoimmunity, which is consistent with the fact that the stress-independent autophagy in TECs is crucial for the thymic self-tolerance. These results suggest that C15ORF48 induces stress-independent autophagy, thereby regulating oxidative stress and self-tolerance.

## INTRODUCTION

Autophagy is a process in which cytoplasmic components of a cell are transported to lysosomes and degraded by multiple enzymes. Initiation of autophagy typically depends on “cellular stress”, such as starvation or mitochondrial damage. Starvation-induced autophagy is often referred to as macroautophagy, which facilitates recycling of key metabolites, such as nucleotides, amino acids, and lipids to support cell survival. Mitochondrial damage-induced autophagy, known as mitophagy, selectively degrades damaged mitochondria and protects cells from mitochondrial stress ^1–4^.

Under normal conditions, the mammalian target of rapamycin (mTOR) protein kinase complex represses autophagy by inactivating Unc-51-like kinase 1 (ULK1) by phosphorylating it at Ser757 ^5,6^. Starvation inhibits mTOR activity, in turn leading to activation of ULK1. ULK1 phosphorylates autophagy-related gene 14 (ATG14), Beclin1, and other autophagy proteins to initiate autophagy ^7^. Mitochondrial stress causes a reduction in intracellular ATP that triggers activation of the AMP-activated protein kinase (AMPK) complex, which is involved in various events related to mitochondrial homeostasis. For instance, if the AMPK complex activates ULK1 by phosphorylating Ser555 ^8^, then, activated ULK1 initiates mitophagy by phosphorylating the key mitophagy-inducing ubiquitin ligase, Parkin ^9^.

In addition to the stress-dependent autophagy, autophagy is induced by some types of stimuli ^10^. IL-6 signaling reportedly triggers macroautophagy in cancer cells ^11^, which may promote chemotherapy resistance in colorectal cancer. Moreover, autophagy is constitutively active in a few tissues such as thymic epithelial cells (TECs), lens epithelial cells, and podocytes, without starvation in mice ^12^. Mechanisms underlying these stress-independent autophagy remain to be elucidated.

TECs are self-antigen-presenting cells and are required for differentiation and selection of major histocompatibility complex (MHC)-restricted and self-tolerant T cells ^13,14^. TECs are separated into cortical TECs (cTECs) and medullary TECs (mTECs), based on their localization in the thymus, and both types of TECs display self-antigen peptide and MHC complexes (self-pMHC) on their cell surfaces. cTECs promote survival of thymocytes expressing T cell antigen receptor (TCR) with moderate affinity for self-pMHC. In contrast, mTECs uniquely express and present thousands of tissue specific self-antigens (TSAs) under regulation of autoimmune regulator (AIRE) and other transcriptional regulators ^15–17^, thereby eliminating T cells that recognize self-pMHC with high affinity or converting them to regulatory T cells (Tregs).

For cell surface expression of self-pMHC by TECs, degradation of cytoplasmic self-proteins is essential for loading self-peptides on surface MHC molecules. Whereas MHC class II normally presents extracellular antigens, autophagy would permit presentation of intracellular antigens on MHC class II through lysosomal pathways. Previous studies revealed that TECs show high autophagy activity without starvation and infections ^12,18^. This implies that such “stress-independent” autophagy might contribute to self-protein degradation for generating self-antigen peptides in TECs. Consistently, some studies have suggested that autophagy in TECs is important for acquisition of T cell self-tolerance ^18–21^. However, the mechanism by which TECs induce autophagy in the absence of starvation is unknown.

C15ORF48 (also known as NMES1, Coxfa4l3, MISTRV, or MOCCI) is a small mitochondrial protein composed of 83 amino acids, which is thought to modulate cytochrome *c* oxidase in electron transport chain complex IV (CIV) ^22–25^. Expression of C15ORF48 is induced by inflammatory stimuli, such as interleukin (IL)-1β, interferon-g, toll-like receptor ligands, and viral infection ^23,24,26^. Nuclear factor-κB (NF-κB) is a pivotal signaling pathway in *C15ORF48* expression ^26^. C15ORF48 reduces CIV activity, mitochondrial membrane potential, and reactive oxygen species (ROS) production, and protects against cell death from viral infection ^23,27^. The 3’ untranslated region (UTR) of *C15ORF48* encodes microRNA miR-147b, which targets the *C15ORF48* homolog, *NDUFA4*. Induction of C15ORF48 is accompanied by miR-147b expression, thereby repressing *NDUFA4* expression via miR-147b ^23,24,26,27^. Previous studies suggested that C15ORF48 replaces NDUFA4 protein in CIV during inflammatory stimuli. This mechanism explains the reduction of CIV activity by expression of C15ORF48 and miR-147b. However, the mechanism underlying the reduction in mitochondrial activity and ROS by C15ORF48 remain unclear ^23^.

In this study, we found that C15ORF48 promotes autophagy independently of starvation or mitochondrial stress. Mechanistically, C15ORF48 reduces mitochondrial activity and intracellular ATP levels, thereby inducing AMPK-ULK1 signaling. Surprisingly, C15ORF48-induced autophagy increased glutathione levels and thereby protected cells from oxidative stress. *C15orf48*^−/–^ mice showed a severe reduction in constitutive autophagy in mTECs and exhibited autoimmunity. Taken together, these results strongly suggest that C15ORF48 is the factor responsible for initiation of “stress-independent” autophagy and that it regulates oxidative stress and self-tolerance.

## RESULTS

### C15ORF48 expression leads to reduction of intracellular ATP and activation of the AMP-activating kinase complex

To specifically investigate the role of C15ORF48 protein, independent of miR-147b, we first established a cell line stably expressing *C15ORF48* lacking its 5′ or 3′ UTR using A549 human lung cancer cells. As previously reported ^22–24,28^, C15ORF48 predominantly localized in mitochondria (Fig. 1a). Forced expression of C15ORF48 reduced mitochondrial membrane potential and intracellular ATP levels (Fig. 1b, c), suggesting that C15ORF48 supresses mitochondrial activity. A reduction in intracellular ATP levels causes activation of AMPK complex ^29^. Consistently, we found that C15ORF48-expressing cells showed increased phosphorylation of AMPKα (Fig. 1d). Thus, these data suggested that increased expression of C15ORF48 reduces mitochondrial activity in the absence of miR-147b, thereby activating AMPK due to the reduction in intracellular ATP.

**Fig. 1:**
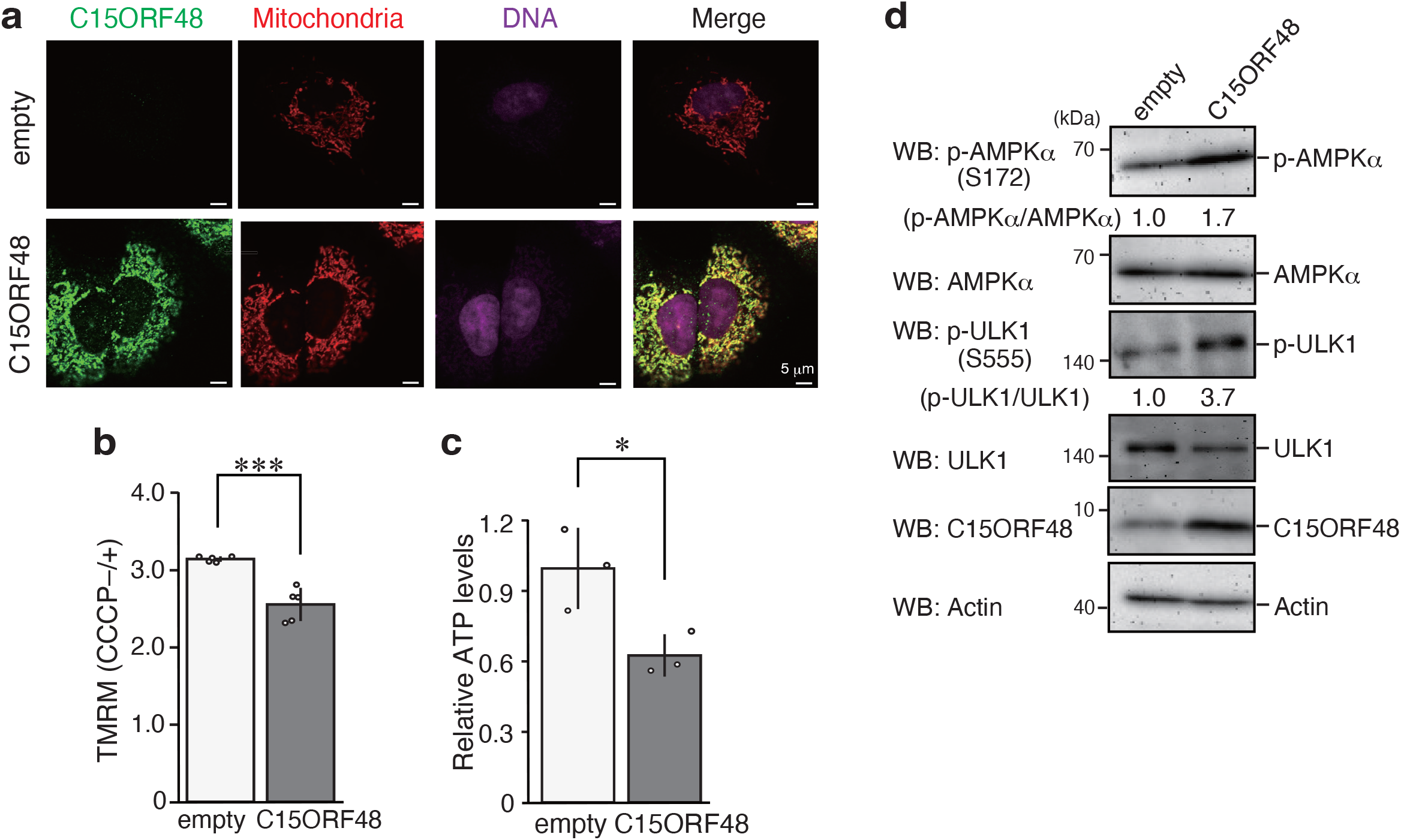
C15ORF48 expression activates AMPK-ULK1 signaling. **a**, A549 cells stably expressing *C15ORF48* (A549/C15ORF48) or the empty (A549/empty) vector were fixed and stained with anti-C15ORF48 antibody and MitoTracker. DNA was counter-stained with TO-PRO-3. **b**, A549/empty and A549/C15ORF48 cells were analyzed for mitochondrial membrane potential. The quantitative ratio of MFI of TMRM in CCCP-untreated (CCCP–) cells to that in CCCP-treated (20 mM, 6 h) cells (CCCP+) was calculated and shown as the mean ± SD. Asterisks indicate statistical significance (***p < 0.001) as calculated using Student’s *t*-test (n = 5). **c**, A549/empty and A549/C15ORF48 cells were analyzed for intracellular ATP levels. Quantitative ratio of ATP concentration normalized to that of controls was calculated and shown as the mean ± SD. Asterisks indicate statistical significance (*p < 0.05) as calculated using Student’s *t*-test (n = 3). **d**, A549/empty and A549/C15ORF48 cells were lysed and subjected to western blotting with indicated antibodies. Band intensity was measured, and quantitative ratios are shown.

### C15ORF48 is an autophagy inducer, independent of starvation

AMPK activates various metabolic processes ^29^, including autophagy ^7,8^. We hypothesized that C15ORF48 could be a natural autophagy inducer because C15ORF48 expression results in phosphorylation of ULK1, a downstream kinase of AMPK, critical for autophagy induction (Fig. 1d), To verify this hypothesis, autophagy induction was evaluated by detecting LC3-II levels and the number of LC3 puncta ascribed to autophagosome formation, hallmarks of autophagy ^30,31^. Detection of autophagosome formation was enhanced by addition of Bafilomycin A1 (Baf A1), which allows sensitive detection of LC3-II accumulation and autophagosome formation by inhibiting fusion of autophagosomes and lysosomes. We found that forced expression of C15ORF48 increased the level of LC3-II and the number of LC3 puncta, even under non-starved conditions, i.e., 4 h after replacement of culture media (Fig. 2a–c). C15ORF48-induced autophagy was repressed by the ULK1 inhibitor, SBI-0206965 (Fig. 2d–f), supporting the involvement of AMPK-ULK1 signaling in C15ORF48-induced autophagy. In contrast, phosphorylated mTOR, a hallmark of starvation-dependent autophagy ^32^, was not reduced by forced expression of C15ORF48 (Fig. 2g), suggesting that starvation-dependent autophagy would not be induced. Thus, C15ORF48-induced autophagy is distinct from the starvation-stress pathway of autophagy.

**Fig. 2:**
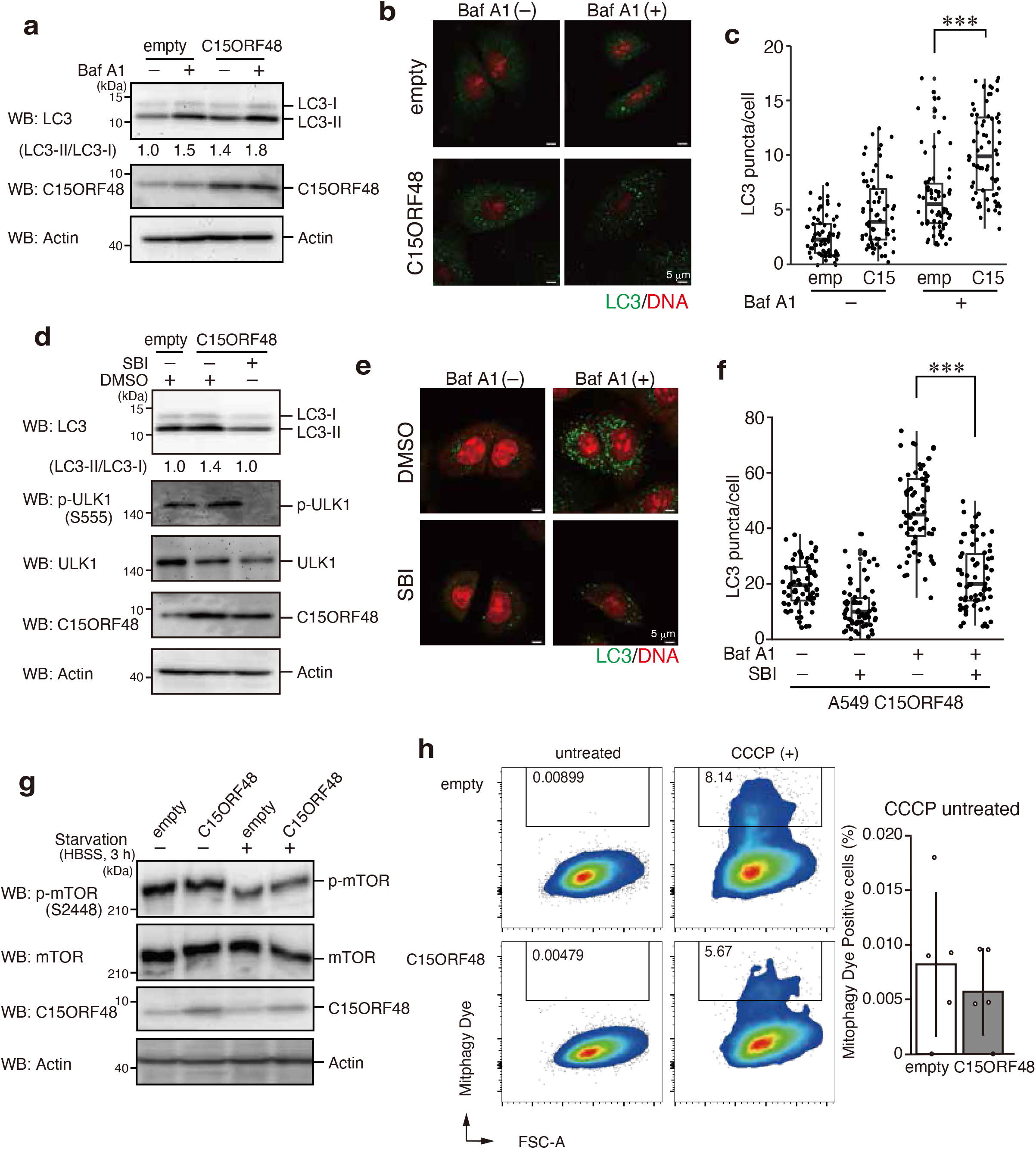
C15ORF48 expression activates autophagy independently of starvation or mitochondrial stress. **a**, A549/empty and A549/C15ORF48 cells were treated with bafilomycin A1 (Baf A1, 200 mM, 1 h) or left untreated 3 h after replacement of culture media. After incubation, cells were lysed and subjected to western blotting as in Fig. 1d. **b**, **c**, A549/empty and A549/C15ORF48 cells were treated with bafilomycin A1 (Baf A1, 200 mM, 1 h) or left untreated 3 h after replacement of culture media. Cells were fixed and stained with anti-LC3 antibodies. DNA was counter-stained with PI (**b**). Numbers of LC3 puncta in each cell were calculated and are shown as means ± SDs. Asterisks indicate statistical significance (***p < 0.001), calculated using Tukey’s multiple comparisons test (n = 70) (**c**). **d**, A549/empty and A549/C15ORF48 cells were incubated for 4 h after replacement of culture media with fresh media containing DMSO (0.1%) or SBI-0206965 (SBI, 10 mM). Whole cell lysates were prepared and subjected to western blotting as in Fig. 1d. **e**,**f**, A549/C15ORF48 cells were treated with bafilomycin A1 (Baf A1, 200 mM, 1 h) or left untreated 3 h after replacement of culture media with fresh media containing DMSO (0.1%) or SBI-0206965 (SBI, 10 mM). Cells were fixed and stained with anti-LC3 antibody. LC3 puncta and DNA were visualized as in Fig. 2b. Numbers of LC3 puncta in each cell were calculated as shown in Fig. 2c. **g**, A549/empty and A549/C15ORF48 cells were unstarved or starved with HBSS (3 h) 1 h after replacement of culture media. Whole cell lysates were prepared and subjected to western blotting with the indicated antibodies. **h**, A549/empty and A549/C15ORF48 cells were treated with Mitophagy dye, and Mitophagy dye-positive cells (boxed areas) were detected by flow cytometry (left). CCCP-treated cells (CCCP+) were analyzed for positive control of mitophagy. Mitophagy dye-positive cells in CCCP-untreated cells (CCCP–) showed basal mitophagy levels. Statistical significance was calculated using Student’s *t*-test (n = 5).

Because C15ORF48 affected mitochondrial activity, C15ORF48-expressing cells might cause mitophagy, leading to selective degradation of mitochondria ^33^. However, the number of mitophagy dye-positive cells did not change in C15ORF48-expressing cells under normal conditions (Fig. 2h) As a control experiment, treatment with the mitochondrial uncoupler, CCCP (carbonyl cyanide *m*-chlorophenylhydrazone), caused mitophagy in C15ORF48-expressing cells, confirming that the machinery for inducing mitophagy was intact. Accordingly, although increased expression of C15ORF48 caused a reduction of mitochondrial activity and AMPK activation, it was not sufficient to induce mitophagy. Overall, these results suggest that C15ORF48 is an inducer of starvation-*independent* macroautophagy, that activates AMPK-ULK1 signaling axis, which is triggered by reduction of intracellular ATP levels due to mild repression of mitochondrial activity.

### C15ORF48 is required for autophagy induced by inflammatory signaling

To determine the role of C15ORF48-induced autophagy in physiological situations, we focused on inflammatory signaling. Inflammatory cytokines, such as IL-1α and tumor necrosis factor (TNF)-α, induced expression of C15ORF48 (Extended Data Fig. 1a, b) ^23,24,26^. These cytokines activate transcription factor NF-κB for expression of its responsive genes. The NF-κB component, RelA, binds to the promoter region of the *C15ORF48* gene, and the NF-κB inhibitor (SC-514) repressed its induction. Thus, NF-κB is necessary for and directly regulates *C15ORF48* mRNA expression (Extended Data Fig. 1c, d).

As expected, C15ORF48 protein up-regulated by IL-1α was predominantly localized in mitochondria (Fig. 3a). Moreover, IL-1α stimulation reduced mitochondrial membrane potential and intracellular ATP levels (Fig. 3b, c). Notably, as observed in forced expression of C15ORF48, IL-1α stimulation increased autophagy activity and activation of AMPK and ULK1, even under non-starved conditions (Fig. 3d–g). Importantly, these IL-1α-inducing responses were repressed by *C15ORF48* knockdown, demonstrating their dependency on C15ORF48 (Fig. 3b, 3c, and g–i). Additionally, the ULK1 inhibitor repressed IL-1α-induced autophagy (Fig. 3j–l). These results suggest that inflammatory stimuli induce expression of C15ORF48 to promotes stress-independent autophagy via AMPK-ULK1 signaling.

**Fig. 3:**
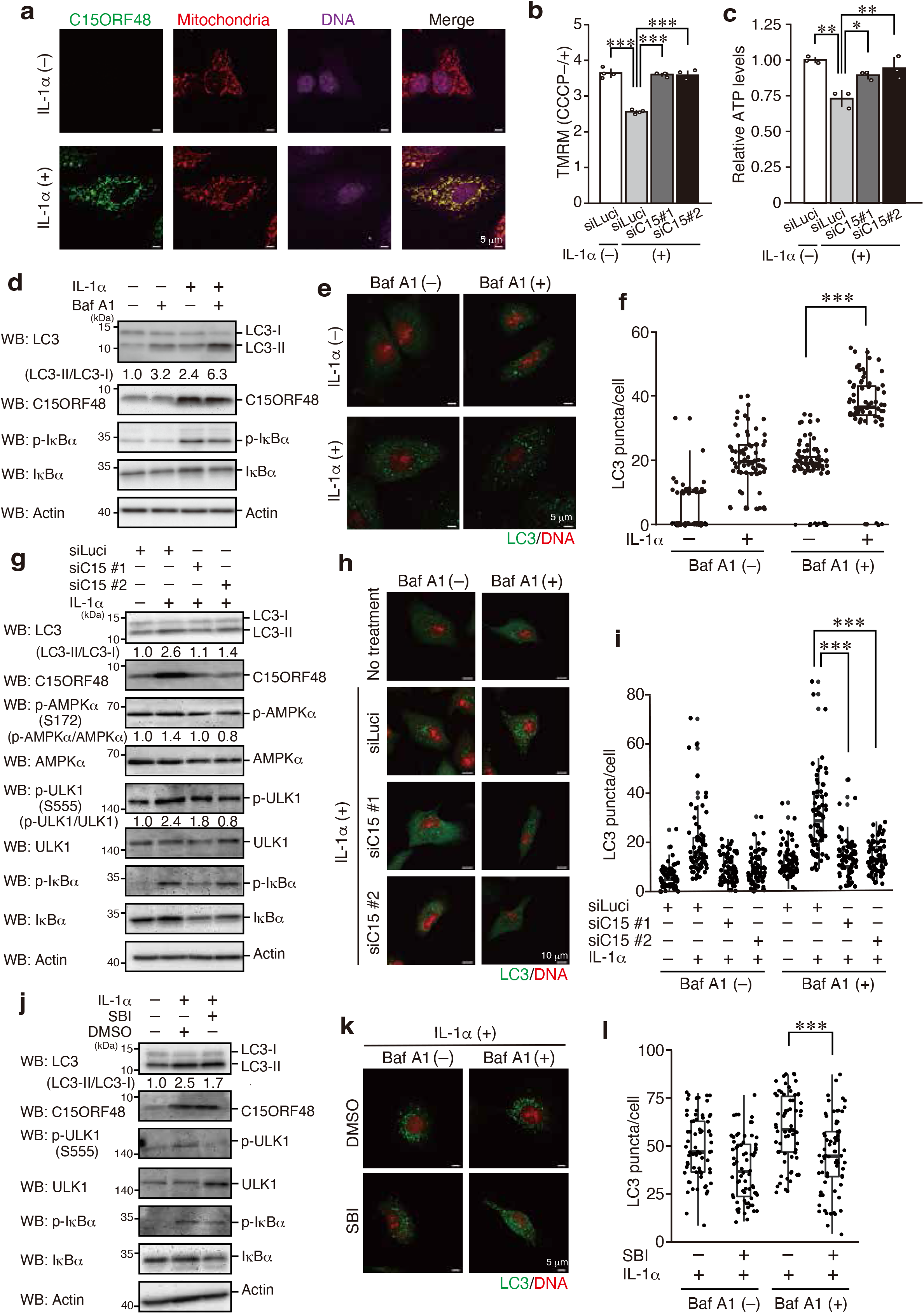
Induction of endogenous C15ORF48 increases autophagy. **a**, A549 cells were stimulated with IL-1α (10 ng/mL) or left unstimulated for 24 h. After incubation, cells were fixed and stained with anti-C15ORF48 antibody and MitoTracker. DNA was counter stained with TO-PRO-3. **b**, A549 cells were transfected with the indicated siRNAs for 48 h and then stimulated with IL-1α (10 ng/mL) or left unstimulated for 24 h. After incubation, cells were treated with or without CCCP (20 mM) for 6 h, followed by TMRM treatment (30 nM) for 30 min. Cells were collected, and MFIs of TMRM were analyzed by flow cytometry. Quantitative ratios of MFI in CCCP-untreated (CCCP–) cells to that in CCCP-treated cells (CCCP+) were calculated and shown as mean ± SD. Asterisks indicate statistical significance (***p < 0.001) as calculated using Tukey’s multiple comparisons test (n = 4). **c**, A549 cells were transfected with the indicated siRNAs for 48 h and then stimulated or left unstimulated with IL-1α (10 ng/mL) for 24 h. After incubation, cells were used for ATP assays. Quantitative ratios of ATP concentration normalized to that of control cells were calculated and shown as means ± SDs. Asterisks indicate statistical significance (**p < 0.01, *p < 0.05) as calculated using Tukey’s multiple comparisons test (n = 3). **d**,**e**,**f**, A549 cells were stimulated or left unstimulated with IL-1α (10 ng/mL) for 20 h. After incubation, media were replaced with fresh media with or without IL-1α for an additional 4 h. Half the samples were treated with bafilomycin A1 (Baf A1, 200 mM) for the final 1 h. Cells were lysed and subjected to western blotting as in Fig. 1d (**d**). Alternatively, cells were fixed and analyzed for LC3 puncta by immunocytochemistry, as in Fig. 2b,c (**e**,**f**). **g**,**h**,**i**, As in Fig. 3d,e,f, except that cells were transfected with the indicated siRNAs for 48 before IL-1α stimulation. **j**,**k**,**l**, A549 cells were stimulated or left unstimulated with IL-1α (10 ng/mL) for 20 h. After incubation, media were replaced with fresh media together with or without IL-1α for an additional 4 h. Some samples were treated with DMSO (0.1%) or SBI-0206965 (SBI, 10 mM) for the final 4 h. Cells were used for western blotting (**j**) and LC3 puncta analysis by immunocytochemistry (**k**,**l**), as in Fig. 3d,e,f.

### High expression of endogenous C15ORF48 increases autophagy

Autophagy promotes survival and metastasis of tumor cells ^34,35^. Therefore, we analyzed highly invasive breast cancer cells, MDA-MB-231, showing constitutively high NF-κB activation ^36^, which might be involved in malignancies ^37^. MDA-MB-231 cells displayed high expression of endogenous C15ORF48 compared with A549 cells, which have lower constitutive NF-κB activity (Fig. 4a). Notably, MDA-MB-231 cells exhibited higher basal autophagy activity than A549 cells (Fig. 4a–c). Furthermore, *C15ORF48* knockdown increased intracellular ATP levels and repressed the phosphorylation levels of AMPKα and ULK1 in MDA-MB-231 cells (Fig. 4d, e). *C15ORF48* knockdown also suppressed basal autophagy activity (Fig. 4e–g). These results suggest that high expression of endogenous C15ORF48 activates stress-independent autophagy in tumor cells.

**Fig. 4:**
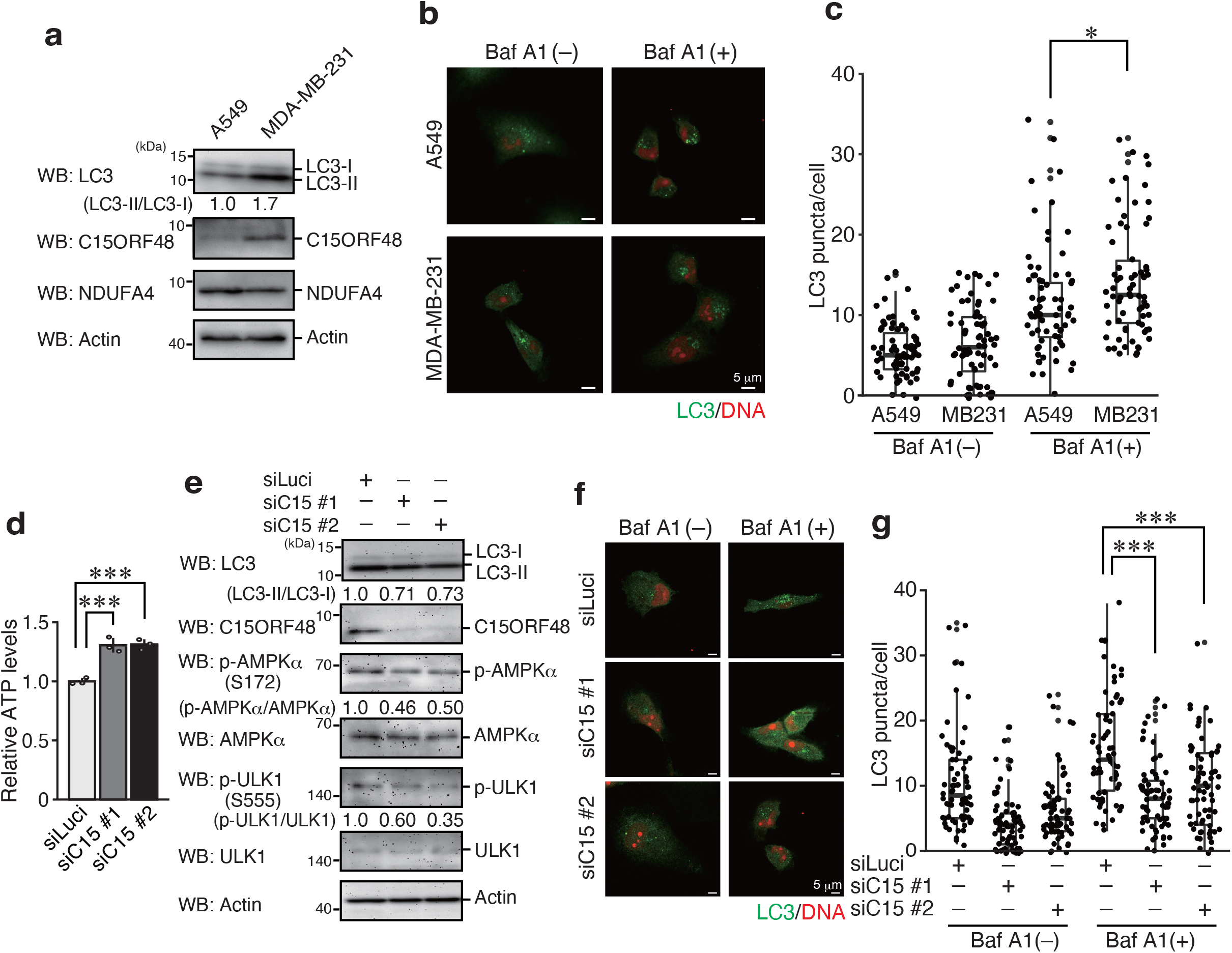
High expression of endogenous C15ORF48 enhances autophagy. **a**, A549 and MDA-MB-231 cells were lysed and subjected to western blotting with the indicated antibodies. **b**,**c**, As in Fig. 2b,c, except that A549 and MDA-MB-231 cells were used. An asterisk indicates statistical significance (*p < 0.05), calculated using Tukey’s multiple comparisons test (n = 70). **d**, MDA-MB-231 cells were transfected with the indicated siRNAs for 48 h. After incubation, cells were used for ATP assays. Quantitative ratios of ATP concentration normalized to that of control cells were calculated and shown as means ± SDs. Asterisks indicate statistical significance (***p < 0.001) as calculated using Tukey’s multiple comparisons test (n = 3). **e**, MDA-MB-231 cells were transfected with the indicated siRNAs for 48 h. 44 h after transfection, media were replaced with fresh media. After incubation, cells were lysed and subjected to western blotting, as in Fig. 1d **f**,**g**, MDA-MB-231 cells were transfected with the indicated siRNAs for 48 h. 44 h after transfection, media were replaced with fresh media. Half the samples were treated with bafilomycin A1 (Baf A1, 200 mM) for the final 1 h. Cells were fixed and analyzed for LC3 puncta by immunocytochemistry, as in Fig 2b,c.

### C15ORF48-induced autophagy increased intracellular glutathione levels to protect cells from oxidative stress

C15ORF48 reduces intracellular ROS levels. However, its mechanism does not appear related to the reduction of CIV activity ^23^. We suspected that C15ORF48 expression could cause an incremental increase in levels of natural antioxidants, thereby reducing ROS levels. Surprisingly, we found that levels of total glutathione and reduced glutathione (GSH) were significantly increased by forced expression of C15ORF48. Importantly, glutathione induction was repressed by *ATG5*- or *ATG7*-knockdown, indicating that autophagy induced by C15ORF48 is responsible for increasing the glutathione level (Fig. 5a–c). Interestingly, forced expression of C15ORF48 protected cells from cell death and repressed caspase-3 activation under oxidative stress condition (H_2_O_2_ treatment) (Fig. 5d, e). In addition, C15ORF48-expressing cells showed increased susceptibility to the glutathione peroxidase (GPX) inhibitor, RSL3 (Fig. 5f). These data suggest that C15ORF48 induces up-regulation of glutathione levels, thereby reducing ROS levels and enhancing resistance to oxidative stress.

**Fig. 5:**
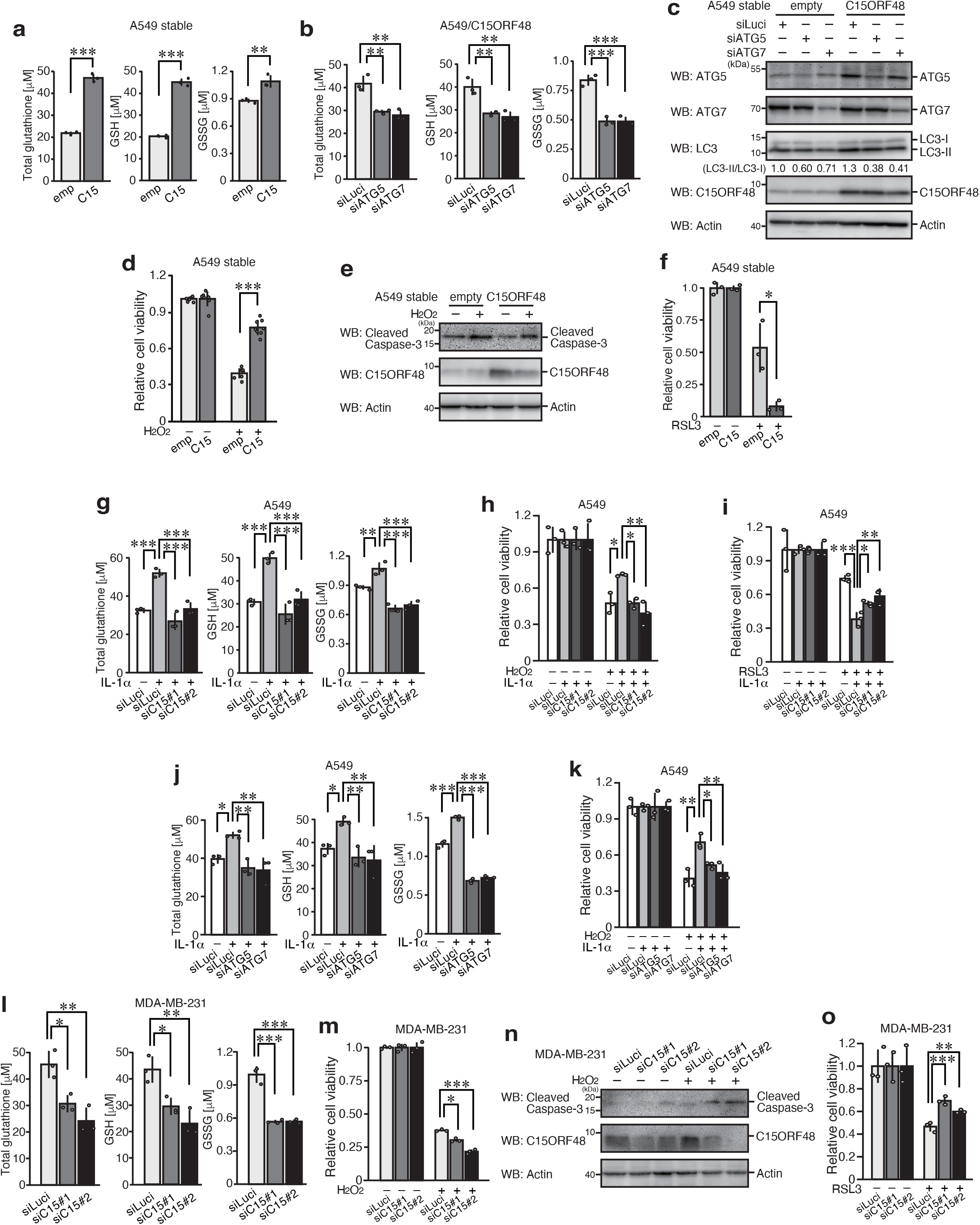
C15ORF48-induced autophagy protects cells from oxidative stress. **a**, A549/empty and A549/C15ORF48 cells were subjected to glutathione assays. Glutathione concentrations were determined and shown as means ± SDs. Asterisks indicate statistical significance (**p < 0.01, ***p < 0.001) as calculated using Student’s *t*-test (n = 3). **b**, A549/C15ORF48 cells were transfected with the indicated siRNAs for 48 h. After incubation, cells were subjected to glutathione assays. Glutathione concentrations were determined and are shown as means ± SDs. Asterisks indicate statistical significance (**p < 0.01, ***p < 0.001), as calculated using Tukey’s multiple comparisons test (n = 3). **c**, A549/empty and A549/C15ORF48 cells were transfected with the indicated siRNAs for 48 h. After incubation, cells were subjected to western blotting, as in Fig. 1d. **d**, A549/empty and A549/C15ORF48 cells were seeded at 20,000 cells/well and treated with or without H_2_O_2_ (300 mM) for 18 h. After incubation, cell viability was calculated with an MTT assay. The quantitative ratio of MTT absorbance in H_2_O_2_-treated cells normalized to that in untreated cells was calculated and is shown as the mean ± SD. Asterisks indicate statistical significance (***p < 0.001) as calculated using Student’s *t*-test (n = 8). **e**, A549/empty and A549/C15ORF48 cells were treated or left untreated with H_2_O_2_ (300 mM) for 6 h. After incubation, cells were subjected to western blotting, as in Fig. 1d. **f**, As in Fig. 5d, except that RSL3 (20 mM, 24 h) was used, instead of H_2_O_2_. Statistical significance (*p < 0.05) was calculated using Student’s *t*-test (n = 3). **g**, A549 cells were transfected with the indicated siRNAs for 24 h. After incubation, cells were treated with with IL-1α (10 ng/mL) or left untreated for 24 h. Then, cells were subjected to glutathione assays, as in Fig. 5b. **h**, A549 cells were seeded at 20,000 cells/well and transfected with the indicated siRNAs for 24 h. After incubation, cells were treated with IL-1α (10 ng/mL) or left untreated for 24 h, and then cells were additionally treated with H_2_O_2_ (300 mM) for 18 h, or left untreated. Cell numbers were calculated by MTT assay, as in Fig. 5d, except that IL-1α-untreated samples were used as control. Asterisks indicate statistical significance (*p < 0.05, **p < 0.01) as calculated using Tukey’s multiple comparisons test (n = 3). **i**, As in Fig. 5h, except that RSL3 (20 mM, 24 h) was used, instead of H_2_O_2_. **j**, As in Fig. 5g. **k**, As in Fig. 5h. **l**, As in Fig. 5b, except that MDA-MB-231 cells were used, instead of A549/C15ORF48 cells. H_2_O_2_ was used at 100 mM for 6 h. Asterisks indicate statistical significance (*p < 0.05, **p < 0.01, ***p < 0.001) as calculated using Tukey’s multiple comparisons test (n = 3). **m**, As in Fig. 5d, except that MDA-MB-231 cells were used instead of A549 stable cells. Asterisks indicate statistical significance (*p < 0.05, ***p < 0.001) as calculated using Tukey’s multiple comparisons test (n = 3). **n**, MDA-MB-231 cells were transfected with the indicated siRNAs for 48 h, and then cells were treated with H_2_O_2_ (100 mM) or left untreated for 6 h. After incubation, cells were subjected to western blotting, as in Fig. 1d. **o**, As in Fig. 5m, except that RSL3 (20 mM, 24 h) was used, instead of H_2_O_2_. Asterisks indicate statistical significance (**p < 0.01, ***p < 0.001) as calculated using Tukey’s multiple comparisons test (n = 3).

Consistently, IL-1α stimulation also increased levels of total glutathione and GSH, and showed enhanced cell viability to oxidative stress and susceptibility to RSL3 (Fig. 5g–i). Importantly, these changes were reversed by *C15ORF48* knockdown (Fig. 5g–i). Moreover, changes in glutathione levels and cell susceptibility to oxidative stress were recovered by *ATG5*- or *ATG7*-knockdown (Fig. 5j, k). Thus, these data suggested that IL-1α stimulation induces C15ORF48-dependent autophagy, which promotes glutathione synthesis, and in turn confers resistance to oxidative stress.

We next tested if the same event occurs in MDA-MB-231 tumor cells expressing high levels of C15ORF48. *C15ORF48* knockdown significantly reduced total glutathione and GSH levels (Fig. 5l). Moreover, *C15ORF48* knockdown depressed resistance to oxidative stress and recovered susceptibility to the GPX inhibitor, RSL3 (Fig. 5m–o). Taken together, these results suggest that C15ORF48 induces stress-independent autophagy and promotes cell survival by eliminating oxidative stress via upregulation of glutathione levels.

### C15ORF48 is involved in development of TECs and mature CD4 single-positive T cells in the thymus

We next sought to determine the physiological function of C15ORF48-inducing autophagy. Because autophagy is active in TECs in the absence of starvation ^12,18^, we speculated that C15ORF48 may be involved in constitutive autophagy induction in TECs. We first investigated expression levels of *C15orf48* mRNA in murine TECs. Analysis of immunological genome project platform data showed that expression of murine *C15orf48* (AA467197) is high in mature mTECs expressing high levels of MHC class II and co-stimulatory CD80 (mTEC^hi^), but low and rare in thymocytes (Extended Data Fig. 2a). Recent single-cell RNA-seq analyses (scRNA-seq) revealed high heterogeneity of TECs. Thus, in addition to separation of cTECs and mTECs, mTECs are further separated based on expression of AIRE and other marker molecules ^38–41^. Re-analysis of scRNA-seq data of murine TECs ^40^ suggested expression of murine *C15orf48* in cTECs (Fig. 6a, cluster 8 and 12), AIRE-expressing (AIRE^+^) mTECs (Fig. 6a, cluster 0 and 2), and a population of post-AIRE mTECs (Fig. 6a, cluster 9, 10, and 14), which are defined as AIRE-negative mTECs, differentiated from AIRE^+^ mTECs. In contrast, *C15orf48* expression is minimal in immature mTECs (Fig. 6a, cluster 3, 4, and 5) and transit-amplifying mTECs (Fig. 6a, cluster 1), which are precursors of AIRE^+^ mTECs ^40,42^. We further performed flow cytometric analysis to confirm C15ORF48 protein expression by using anti-murine C15ORF48 antibody. Because of relatively high non-specific binding of this antibody in flow cytometric analysis, the mean fluorescent intensity (MFI) for antibody binding in wild-type TECs was subtracted from that in TECs from *C15orf48*-deficient (*C15orf48*^−/–^) mice (Extended Data Fig. 2b, c). Importantly, whereas C15ORF48 protein is abolished by the CRISPR-Cas9 system in these mutant mice, the expression level of NDUFA4 protein, which is regulated by miR-147b encoded in the 3′-UTR of the *C15orf48* gene ^23,24,26,27^, was not affected (Extended Data Fig. 2d). TECs (EpCAM^+^CD45^−^ TER119^−^ cells) were separated into cTECs and mTECs, based on expression of Ly51 and binding of UEA-1 lectin (Fig. 6b). mTECs (UEA-1^+^Ly51^−^) were further divided into mTECs^lo^ and mTECs^hi^ by the expression level of CD80 (Fig. 6b). C15ORF48 protein expression was detected in mTECs^hi^ and cTECs, but not mTECs^lo^. mTECs^hi^ were further sub-classified into Early-, Late-, and Post-Aire mTECs based upon CD24 and Sca1 expression ^43^. Flow cytometric analysis showed C15ORF48 protein expression in all of these subpopulations (Fig. 6c). Consequently, C15ORF48 is expressed in mature types of TECs, implying its functions in self-antigen presentation through autophagy-dependent self-protein degradation.

**Fig. 6:**
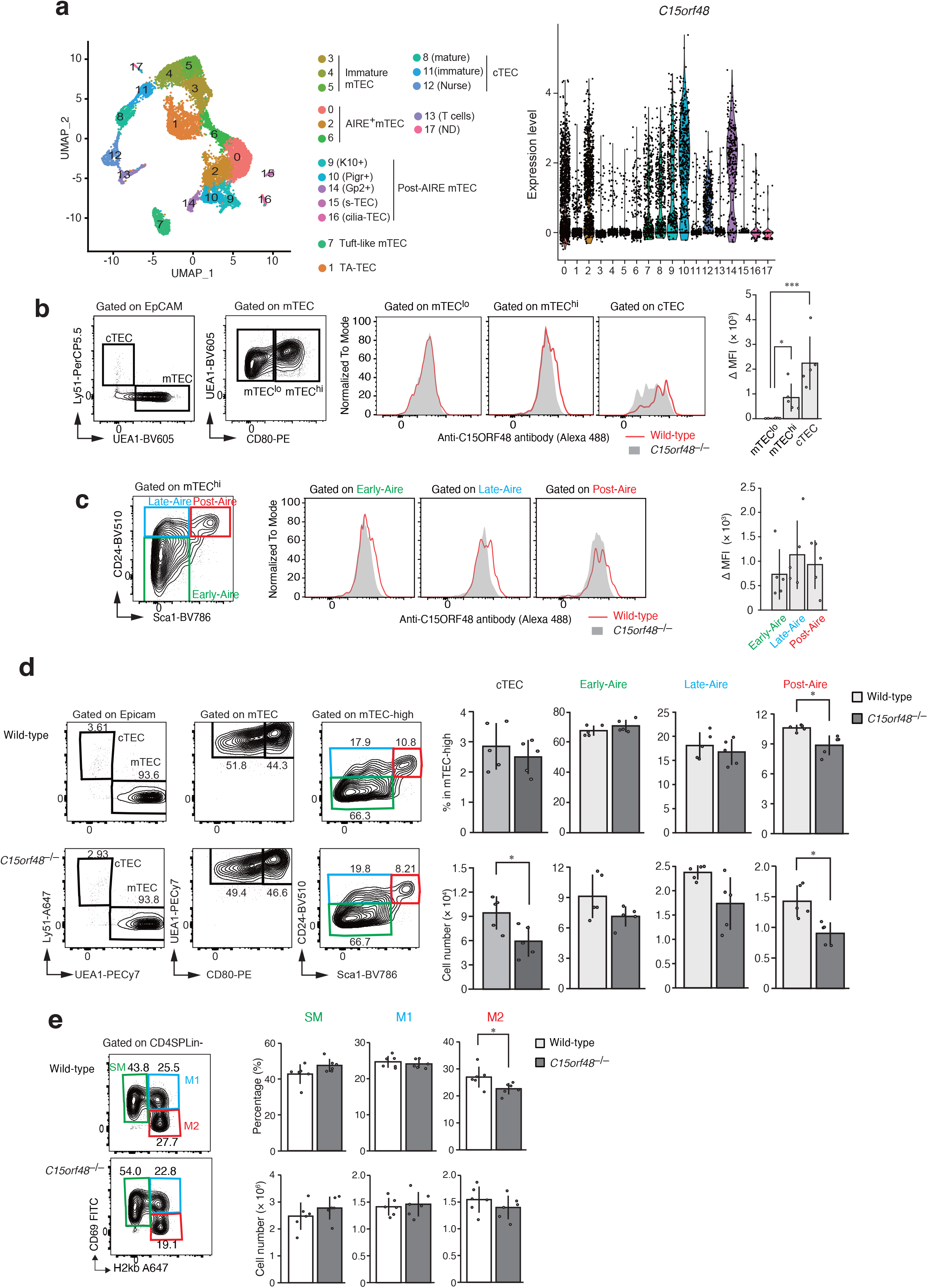
C15ORF48 is expressed in cTECs and mature mTECs and is involved in development of TECs and mature CD4 single-positive T cells in the thymus. **a**, Uniform manifold approximation and production (UMAP) plot of scRNA-seq data from TECs (EpCAM^+^ CD45^−^ TER119^−^) from 4-week-old mice. Cell clusters (R0 to R17) are indicated by colors and numbers in the plot (left). Violin plots depicting expression levels of *C15orf48* in each cluster (right). **b**, Representative images of flow cytometry plots of cTECs, mTEC^lo^ and mTEC^hi^ in wild-type mice (left). Representative images of MFIs of anti-C15orf48 in mTEC^lo^, mTEC^hi^, and cTEC from wild-type and *C15orf48*^−/–^ mice (middle). MFIs of anti-C15orf48 in TECs from wild-type mice were subtracted from those in TECs from *C15orf48*^−/–^ mice and are shown as means ± SDs. Asterisks indicate statistical significance (*p < 0.05, ***p < 0.001), as calculated using Tukey’s multiple comparison test (n = 5) (right). **c**, As in Fig. 6b, except that data of Early-Aire, Late-Aire, and Post-Aire mTEC subfractions are shown, instead of those of cTECs, mTEC^lo^ and mTEC^hi^. **d**, Representative images of flow cytometry plots of cTECs, mTEC^lo^, mTEC^hi^ and the three mTEC subfractions in wild-type and *C15orf48*^−/–^ mice (left). Numbers of cTECs, Early-Aire, Late-Aire, and Post-Aire mTECs and their ratios to total thymic cells are shown as means ± SDs. Asterisks indicate statistical significance (*p < 0.05), as calculated using Student’s *t*-test (n = 5) (right). **e**, Representative images of flow cytometry plots of SM, M1, and M2 CD4SP cells in wild-type and *C15orf48*^−/–^ mice (left). Numbers of SM, M1, and M2 CD4SP cells and their ratios to total thymocytes are shown as means ± SDs. Asterisks indicate statistical significance (*p < 0.05), as calculated using Student’s *t*-test (n = 5) (right).

We next investigated influences of C15ORF48 depletion in thymic cell development. The weight of the thymus and total thymic cell number did not differ significantly between wild-type and *C15orf48*^−/–^ mice (Extended Data Fig. 3a). Numbers of cTECs and Post-Aire mTECs were slightly reduced in *C15orf48*^−/–^ mice (Fig. 6d). This suggests that C15ORF48 participates in maintenance of cTECs and Post-Aire mTECs, which suggests a function of C15ORF48 in cell survival.

In the thymus, CD4 and CD8 single-positive thymocytes (CD4SP and CD8SP) differentiate from bone marrow-derived progenitors via the stage of CD4 and CD8 double-positive thymocytes (DP) ^44,45^. Flow cytometric analysis using CD4 and CD8 antibodies suggested that numbers and ratios of each subset to total thymocytes were not significantly affected in *C15orf48*^−/–^ mice (Extended Data Fig. 3b). CD4SPs and CD8SPs in the thymic medulla are further subdivided into three subpopulations, according to their maturation levels: semi-mature (SM), mature 1 (M1), and mature 2 (M2) ^46^. Intriguingly, the ratio of the M2 subfraction in CD4SP cells was slightly reduced in *C15orf48*^−/–^ mice (Fig. 6e) whereas no apparent change was detected in the number or ratio of CD8SP cells (Extended Data Fig. 3c). Thus, C15ORF48 may contribute to development or selection of mature CD4SP cells, likely through TEC functions.

### C15ORF48 is critical for stress-independent autophagy induction in TECs

We addressed the requirement of C15ORF48 for stress-independent autophagy in TECs. *C15orf48*^−/–^ mice were crossed with mice expressing green fluorescent protein (GFP)-labeled LC3 (GFP-LC3) to monitor autophagosome formation ^12^. As reported previously, GFP-LC3 puncta were detected in the thymus of control mice without starvation (Fig. 7a). Co-immunostaining showed that GFP puncta are formed in cells expressing Keratin-5, confirming high autophagy activity in TECs without starvation (Fig. 7a). Strikingly, the number of GFP-LC3 puncta in TECs is significantly diminished in the absence of C15ORF48 (Fig. 7a). In contrast, GFP-LC3 puncta emerging during fasting are comparable in skeletal muscles between controls and *C15orf48*^−/–^ mice, suggesting that starvation-induced autophagy is not affected by depletion of C15ORF48 (Fig. 7b). Taken together, these results indicate that C15ORF48 controls stress-independent autophagy in TECs, but not in typical starvation-induced autophagy. Because autophagy in TECs contributes to T cell self-tolerance, we wondered whether *C15orf48*^−/–^ mice exhibit signs of autoimmunity. Although no apparent changes were observed in cell numbers or ratios of effector memory T cells or Tregs in lymph node T cells of *C15orf48*^−/–^ mice (Extended Data Fig. 4a–e), sera from *C15orf48*^−/–^ mice showed high immune reactivities against mouse tissue sections, such as lacrimal glands, submandibular glands, eyes, and ovaries (Fig. 7c), suggesting that sera from *C15orf48*^−/–^ mice contain autoantibodies reactive to self-antigens in these organs. Overall, these results suggest that C15ORF48 contributes to establishing self-tolerance.

**Fig. 7:**
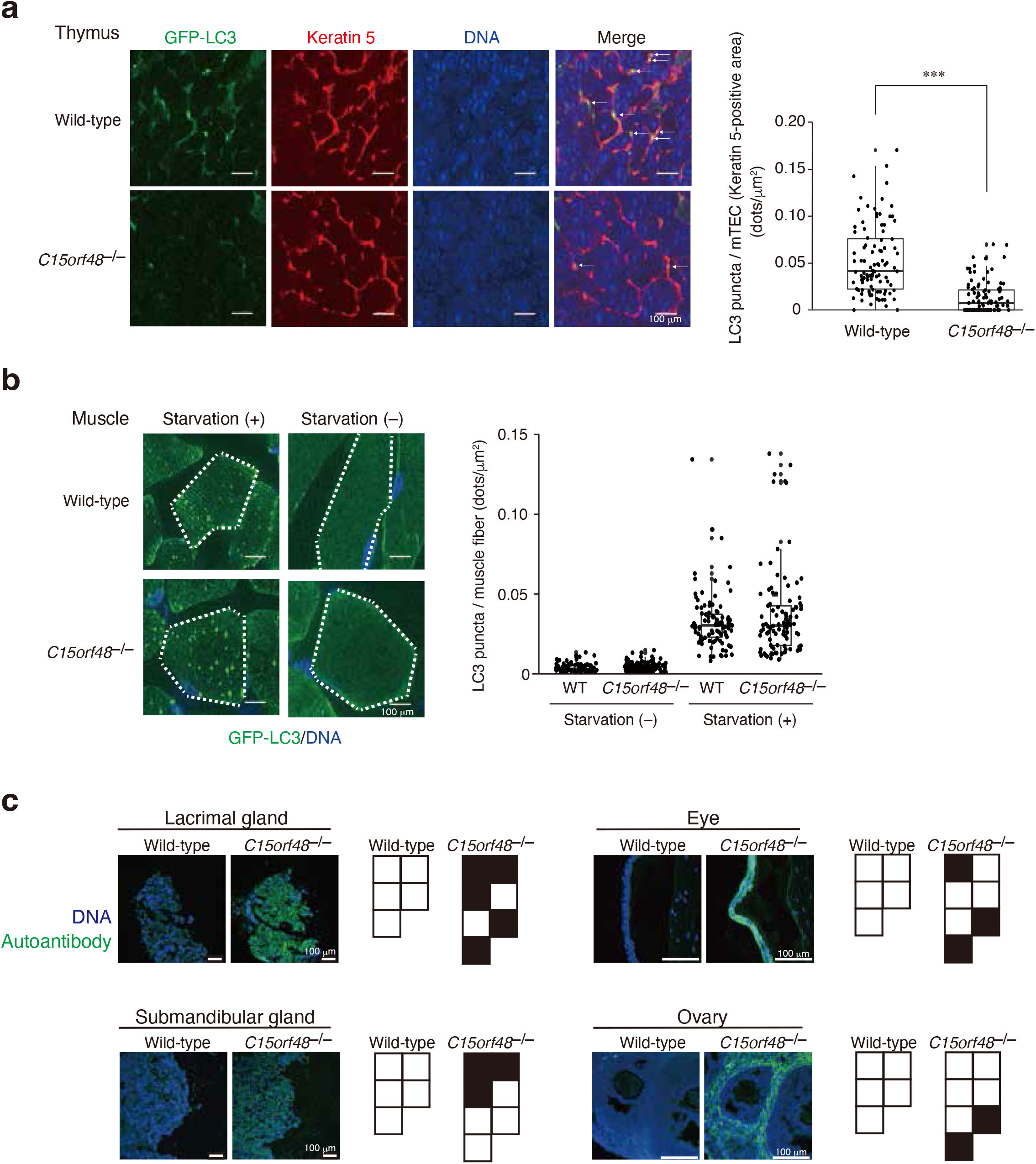
C15ORF48 is critical for stress-independent autophagy in TECs. **a**, Thymus sections from normal fed wild-type and *C15orf48*^−/–^ mice were stained with anti-GFP and anti-Keratin 5 antibodies and DAPI (left). GFP-LC3 puncta in Keratin 5-positive areas are indicated by arrows (left panel). Numbers of GFP-LC3 puncta in Keratin 5-positive areas are shown (right panel). Asterisks indicate statistical significance (***p < 0.001), as calculated using Student’s *t*-test (n = 100). **b**, GFP-LC3 mice (wild-type) and GFP-LC3/*C15orf48*^−/–^ (*C15orf48*^−/–^) mice were starved for 48 h. After fasting, thigh skeletal muscle sections were prepared and stained with an anti-GFP antibody and DAPI. Single muscle fibers are encircled by broken lines (left panel). The number of GFP-LC3 puncta in the muscle area is shown (right panel). Statistical significance was calculated using Student’s *t*-test (n = 100). Wild-type, WT. **c**, Immunostaining of tissue sections from *Rag1*^−/–^ mice with sera from 21-week-old wild-type and *C15orf48*^−/–^ mice. Nuclei were counter-stained with DAPI. Each box represents serum from a single mouse (wild-type, n = 5; *C15orf48*^−/–^, n = 7). The black box indicates the detection of reactivity in tissue sections from *Rag1*^−/–^ mice.

## DISCUSSION

In this study, we found that the mitochondrial protein, C15ORF48, activates autophagy (Extended Data Fig. 5). Autophagy is generally accepted as a phenomenon induced by cellular stress, such as starvation or mitochondrial damage, and prevents cell death by promoting recycling of nutrients or eliminating damaged organelles. In contrast to stress-dependent autophagy, C15ORF48-induced autophagy is “independent of cellular stress” for the following three reasons: (1) autophagy is activated even in non-starved conditions, confirming that mTOR phosphorylation was not reduced by C15ORF48 expression, (2) mitophagy is not activated in C15ORF48-expressing cells, and (3) C15ORF48 is required for constitutive autophagy in TECs and is dispensable in starvation-dependent autophagy in murine skeletal muscle.

Our results show that C15ORF48 reduces mitochondrial membrane potential and lowers intracellular ATP levels, thereby leading to autophagy via AMPK-ULK1 signaling. Repression of mitochondrial membrane potential causes mitochondrial stress and subsequent mitophagy ^47,48^, whereas mitophagy did not increase in C15ORF48-expressing cells. Therefore, C15ORF48 is likely to alleviate, rather than exacerbate, mitochondrial activation and to promote cell survival by inducing stress-independent autophagy.

Furthermore, our present data show that C15ORF48-induced autophagy increases intracellular glutathione levels and prevents oxidative stress. Glutathione is a tripeptide composed of three amino acids (glutamic acid, cysteine, and glycine) and acts as an important cofactor for the antioxidant enzyme, GPX ^49^. C15ORF48-expressing cells showed higher susceptibility to a GPX inhibitor, suggesting that C15ORF48-expressing cells strengthen the glutathione-GPX antioxidant system for survival. Given that nonselective macroautophagy is involved in recycling of nutrients ^1,3^, C15ORF48-induced autophagy may be classified as nonselective macroautophagy that facilitates degradation of proteins and recycling of amino acids for glutathione synthesis.

In the thymus, *C15orf48* is expressed in cTECs and mTECs, but its regulation in these cells remains unclear. It is known that NF-κB signaling is pivotal in TEC differentiation. Differentiation of mTECs is controlled by the TNF receptor superfamily member receptor activator of NF-κB (RANK) and CD40 ^50–52^, the NF-κB component RelB ^53^, NF-κB-inducing kinase (NIK) ^54^, and NF-κB signal transducer TNF receptor-associated factor 6 (TRAF6) ^55^. Mice harboring mutations of these genes show defects in mTEC differentiation. NIK is also involved in cTEC differentiation ^56^. Given that NF-κB is a key transcription factor for *C15ORF48* expression in human cancer cell lines, NF-κB signaling is likely to be involved in *C15orf48* expression in cTECs and mTECs.

In the present study, *C15orf48*^−/–^ mice showed reduced autophagic activity in mTECs. Furthermore, *C15orf48*^−/–^ mice showed a reduction in the M2 subfraction of CD4SP cells and the increment of autoantibodies in sera. Autophagy in mTECs is important for negative selection of CD4SP cells because autophagy substrates are loaded on MHC class II and preferentially presented to CD4SP thymocytes as self-antigens ^18,19,57^. Therefore, expression of C15ORF48 is likely to trigger autophagy for digestion and presentation of TSAs in mTECs. In addition, *C15orf48*^−/–^ mice showed a slight reduction in cTECs and mTECs (Post-Aire), suggesting that C15ORF48-induced autophagy is involved in maintenance of these cells via suppression of basal oxidative stress. Therefore, it is possible that a reduction of TECs affects differentiation and selection of CD4SP cells in *C15orf48*^−/–^ mice.

Recent studies have shown that H_2_O_2_-mediated redox status is closely associated with basal autophagy and negative selection of thymocytes in TECs using transgenic mice overexpressing the hydrogen peroxide quenching enzyme, catalase ^21^. Although the mechanism by which H_2_O_2_ regulates autophagy in TECs is unclear, AMPK-ULK1 signaling is a candidate target of H_2_O_2_ in controlling autophagy ^58^. Because AMPK-ULK1 signaling is activated downstream of C15ORF48, this signaling is likely to be a key mediator of constitutive autophagy in TECs.

Cancer cells exhibit constitutive autophagy, which enhances their survival and proliferation ^10^. However, molecular mechanisms underlying starvation-independent autophagy in cancer cells remain largely unknown. In this study, we showed that human breast cancer (MDA-MB-231) cells have high expression of C15ORF48 and high basal autophagy activity. Notably, C15ORF48 increases glutathione levels and reduces oxidative stress in these cells. MDA-MB-231 cells have constitutive activation of NF-κB ^36^, which is frequently observed in various types of cancer cells and is important in cancer progression. Because constitutively active NF-κB in cancer cells drives *C15ORF48* expression ^59^, C15ORF48 may be pivotal in constitutive autophagy and may promote cancer cell survival by preventing basal oxidative stress.

In this study, we found that *C15orf48*^−/–^ mice show reduced stress-independent autophagy in mTECs and exhibit autoimmunity, suggesting that C15ORF48 is involved in negative selection of thymocytes and acquisition of self-tolerance. *C15orf48*^−/–^ mice have autoantibodies against eyes and ovaries, which were also observed in *Aire*-deficient mice ^15^. These observations support the notion that C15ORF48 is involved in AIRE-dependent negative selection of thymocytes. *AIRE* was identified as the primary causative gene in human patients suffering from an autoimmune disease ^60–62^. Autoantibodies against lacrimal glands, submandibular gland, and eyes, which were detected in *C15orf48*^−/–^ mice, are pathological characteristics of autoimmune diseases, such as Sjögren’s syndrome ^63–65^. Therefore, it is possible that inactivation of the *C15ORF48* gene causes human autoimmune diseases. Mutations and polymorphisms in the *C15ORF48* gene in autoimmune disease patients should be analyzed in the future.

## ACKNOWLEDGMENTS

The authors thank Dr. Noboru Mizushima for GFP-LC3 mice, Dr. Yasuyuki Kurihara, Dr. Kazumasa Aoyama, Mr. Takuro Araki, and Mr. Naoto Hori for their technical assistance, and members of the central facilities of RIKEN IMS for generation of *C15orf48*^−/–^ mice. This work was supported in part by grants-in-aid for Challenging Research (Exploratory) [grant number 19K22482 (to Noritaka Y)], Scientific Research (C) [grant numbers 21K08121 (to HT) and 21K06543 (to Noritaka Y)], and Research Activity Start-up [grant number 22K20706 (to YT)] from the Japanese Ministry of Education, Culture, Sports, Science, and Technology, Chiba Foundation for Health Promotion & Disease Prevention (to YT), Takeda Science Foundation (to Noritaka Y), and the Research Foundation for Pharmaceutical Sciences (to Noritaka Y), Grants-in-Aid for Scientific Research from JSPS (20K07332, 20H03441) (TA, NA), and CREST from Japan Science and Technology Agency (JPMJCR2011) (TA and TJK).

## AUTHOR CONTRIBUTIONS

YT, TA and Noritaka Y designed the study, performed experiments, analyzed results and wrote the manuscript. Moeka M, NT, YK, SK, KH, Maki M, TI, NA, TS, TM, MH, RE, HI, YM, NH, TJK, Naoto Y, and HT assisted with experiments and provided several reagents. TA and Noritaka Y supervised and directed the research. All authors discussed the results and commented on the manuscript.

## FIGURE LEGENDS

**Extended Data Fig. 1: Induction of C15ORF48 expression by NF-κB signaling.**

**a**, A549 cells were stimulated with IL-1α (10 ng/mL) or TNF-a (10 ng/mL) for the indicated times. After incubation, cells were analyzed for *C15ORF48* mRNA expression by qPCR. *GAPDH* mRNA expression was used to normalize these data. The expression level of *C15ORF48* mRNA in unstimulated cells was set to 1. Results represent the mean ± SD (n =3). Asterisks indicate statistical significance (***p < 0.001) as calculated using Tukey’s multiple comparisons test (n = 3).

**b**, A549 cells were stimulated with IL-1α (10 ng/mL) or TNF-a (10 ng/mL) for the indicated times. After incubation, cells were subjected to western blotting with the indicated antibodies.

**c**, Schematic diagram of *C15ORF48* promoter region (upper). The region targeted by the designed primer pair is shown (Amplicon). TSS, transcription start site. A549 cells were stimulated with IL-1α (10 ng/mL) for the indicated times. After incubation, cells were analyzed by ChIP assays with anti-RelA or control antibodies to evaluate the binding activity of RelA to the *C15ORF48* promoter region (lower). Percentage input values are shown relative to those of control immunoprecipitates. These results represent the mean ± SD. Asterisks indicate statistical significance (**p < 0.01) calculated using Student’s *t*-test (n = 3).

**d**, A549 cells were stimulated with IL-1α (10 ng/mL) or left untreated for 24 h. 21 h after stimulation, cells were treated with the NF-κB inhibitor SC514 (100 mM) or without. Some samples were further treated with bafilomycin A1 (Baf A1, 200 mM) for the final 1 h. After incubation, cells were subjected to western blotting with the indicated antibodies.

**Extended Data Fig. 2: A 7-bp deletion in the exon 3 of *C15orf48* gene in *C15orf48*^−/–^ mice.**

**a**, Expression levels of murine *C15orf48* (*AA467197*) in thymic cells in immunological genome project platform data.

**b**, Sequencing data of CRISPR-Cas9-targeted regions in *C15orf48* exon 3 from wild-type and *C15orf48*^−/–^ mice. The boxed sequence indicates a 7-bp deletion in *C15orf48*^−/–^ mice (upper). The deleted region in *C15orf48*^−/–^ mice is shown as a gray bar. *C15orf48* genome sequences and corresponding amino acid sequences of wild-type and *C15orf48*^−/–^ mice are shown (lower).

**c**, Schematic diagram of *C15orf48* genomic structure. Primer F1 recognizes the region corresponding to the 7-bp deletion in *C15orf48*^−/–^ mice; therefore, the primer pair between F1 and R cannot amplify PCR products from *C15orf48*^−/–^ mouse genomic DNA (upper). A representative image of genotyping PCR of wild-type and *C15orf48*^−/–^ mice using three primers (Primer F1, F2, and R) (lower).

**d**, Extracts from testes of wild-type and *C15orf48*^−/–^ mice were subjected to western blotting with the indicated antibodies.

**Extended Data Fig. 3: Development of thymocytes in wild-type and C15orf48–/– mice.**

**a**, Ratios of thymus weight to whole body weight (left), total cell numbers in thymus (middle), and ratios of TECs to whole thymic cells (right) in wild-type and *C15orf48*^−/–^ mice are shown as means ±

SDs. Statistical significance was calculated using Student’s *t*-test (n = 5).

**b**, Representative images of flow cytometry plots of DN, DP, CD4SP, and CD8SP thymocytes in wild-type and *C15orf48*^−/–^ mice (left). Numbers of DN, DP, CD4SP, and CD8SP cells and their ratios to total thymocytes are shown as means ± SDs. Statistical significance was calculated using Student’s *t*-test (n = 3) (right).

**c**, Representative images of flow cytometry plots of SM, M1, and M2 CD8SP cells in wild-type and *C15orf48*^−/–^ mice (left). Numbers of SM, M1, and M2 CD8SP cells and their ratios to total thymocytes are shown as means ± SDs (n = 7) (right).

**Extended Data Fig. 4: Memory T cells and Tregs in secondary lymph nodes of wild-type and C15orf48–/– mice.**

**a,b**, Weights (**a**) and total cell numbers (**b**) of brachial lymph nodes (BL) and inguinal lymph nodes (IL) and in 21-week-old wild-type and *C15orf48*^−/–^ mice are shown as means ± SDs. Statistical significance was calculated using Student’s *t*-test (n = 3).

**c**, Representative images of flow cytometry plots of memory CD4 and CD8 T cells in BL in wild-type and *C15orf48*^−/–^ mice. Total numbers of memory CD4 and CD8 T cells and ratios of these cells to total cell numbers in BL are shown as means ± SDs. Statistical significance was calculated using Student’s *t*-test (n = 3).

**d**, As in Extended Data Fig. 3c, except that memory CD4 and CD8 T cells in IL were analyzed.

**e**, As in Extended Data Fig. 3c, except that Tregs in BL and IL were analyzed.

**Extended Data Fig. 5: A schematic model of C15ORF48-induced, stress-independent autophagy.**

The mitochondrial protein, C15ORF48, a subunit of electron transport chain complex IV, is highly expressed in several cancer cells and thymic epithelial cells, and NF-kB signaling is important for its expression. C15ORF48 reduces mitochondrial activity and intracellular ATP levels, thereby inducing autophagy via pro-autophagic AMPK-ULK1 signaling independently of starvation or mitochondrial stress. C15ORF48-induced autophagy in cancer cells increases intracellular glutathione levels and prevents oxidative stress. C15ORF48-induced autophagy in thymic epithelial cells regulates self-tolerance and prevents autoimmunity.

## METHODS

### Cell culture and antibodies

A549 and MDA-MB-231 cell lines were purchased from ATCC and cultured in Dulbecco’s modified Eagle’s medium (DMEM) supplemented with 5% fetal bovine serum at 37°C. To establish stable cell pools, A549 cells were transfected with the pIRESpuro3-CAG/C15orf48 vector or an empty vector using Lipofectamine2000 (Thermo Fisher Scientific) following the manufacturer’s instructions. Transfected cells were selected for puromycin resistance. siRNAs (5 nM) were transfected using the reverse transfection method with Lipofectamine RNAiMax (Thermo Fisher Scientific), according to the manufacturer’s instructions. Silencer select siRNAs specific to *C15ORF48* #1 (s38981), *C15ORF48* #2 (s228367), *ATG5* (s18160), and *ATG7* (s20650) were purchased from Thermo Fisher Scientific. siRNA for luciferase (sense:5′-GCGCUGCUGGUGCCAACCCTT-3′ and antisense:5′-GGGUUGGCACCAGCAGCGCTT-3′) was purchased from Hokkaido System Sciences and used as a control siRNA. Recombinant human IL-1α and TNF-α was purchased from PeproTech. The IKK inhibitor SC-514 was purchased from Cayman Chemical. Antibodies used in this study are shown in Supplementary Table S1.

### Mouse models

All mice were maintained under pathogen-free conditions and handled in accordance with Guidelines of the Institutional Animal Care and Use Committee of RIKEN, Yokohama Branch (2018-075). Almost all available mutant and control mice were randomly used for experiments without selection.

C57BL/6 mice were purchased from CLEA. Littermates and age-matched wild-type mice were used as controls. GFP-LC3 mice (RBRC00806) were obtained from RIKEN BRC. C57BL/6 *Rag1*^−/–^ mice were obtained from the Department of Animal Management at RIKEN.

*C15orf48* knockout mice were generated using the CRISPR-Cas9 system with fertilized eggs derived from C57BL/6J mice. C57BL/6J female mice were injected with 5 IU Pregnant Mare Serum Gonadotropin (PMS). After 48 h, the mice were injected with 5 IU human chorionic gonadotropin (hCG). Serially injected female mice were mated with C57BL/6J male mice. Fertilized eggs were isolated from mating plugs of mated female mice. Synthetic crRNA (Alt-R CRISPR-Cas9 crRNA, IDT, 16 ng/mL), tracrRNA (Alt-R CRISPR-Cas9 tracrRNA, IDT, 24 ng/mL), and Cas9 nuclease (Alt-R S.p. HiFi Cas9 Nuclease V3, 100 ng) were dissolved in 100 mL of Opti-MEM I and electroporated into the embryos. The sequence of the crRNA was as follows: 5′-ACGCUUAUAAAAAACGCCAAGUUUUAGAGCUAUGCU-3′. Electroporation was performed as described previously ^66^. To obtain newborn mice, 2-cell stage embryos were transferred to oviducts of pseudopregnant ICR females.

Genomic DNA of offspring was extracted from tail biopsies using 50 mM NaOH and incubated for 20 min at 95□. Extracted samples were chilled on ice and then mixed with 1M Tris-HCl (pH 8.0). After mixing, samples were centrifuged at 4□ and collected supernatant was used for genotyping PCR. The following three primers were used for PCR: 5′-CTCAGCTCATTCCTTTGGCG-3′ (sense, F1), 5′-GGTCCGGGGGAAGATGTTTT-3′ (sense, F2), and 5′-GGCTCGAACTGTGTAGGCAT-3′ (antisense, R). After initial denaturation at 94°C for 2.5 min, PCR was performed for 30 cycles (15 s at 94°C, 15 s at 55°C, and 30 s at 72°C), followed by 5 min additional extension at 72°C using the BIOTAQ^TM^ DNA polymerase (NIPPON Genetics). While 618-bp DNA fragments were amplified from the wild-type genomic DNA with primer pairs F1 and R, 836-bp DNA fragments were amplified from *C15orf48*^−/–^ genomic DNA with primer pairs F2 and R.

### Plasmids

The cDNA fragment encoding C15orf48 (lacking the 5′ or 3′ UTR) was generated by PCR from the reverse-transcribed product of A549 cells total RNA. The following primers were used for PCR:5′-TAGCTTAAGCCACCATGAGCTTTTTCCAACTCCTGATGAAAAGG-3′ (sense) and 5′-TCAGAATTCTCATTTGGTCACCCTTTGGACATTTTGCAA-3′ (antisense). The synthesized cDNA fragment was inserted into the AflII and EcoRI sites of the pIRESpuro3-CAG vector ^67^.

### Western blotting

Cells were washed with phosphate-buffered saline (PBS), directly lysed with Laemmli buffer, and used as whole-cell lysates. Testes from 4-week-old male mice and tissues from *Rag1*^−/–^ female mice were washed with PBS and then homogenized in TNE buffer (10 mM Tris-HCl, 150 mM NaCl, 10 mM Triton X-100, and cOmplete^TM^ (Sigma-Aldrich); pH 7.4). Homogenized tissues were sonicated and then centrifuged at 10,000 × *g* for 5 min at 4□. Supernatants were collected and mixed with Laemmli buffer after measuring the protein concentration using a TaKaRa BCA Protein Assay Kit (TaKaRa). Samples were subjected to SDS-polyacrylamide gel electrophoresis and electrotransferred onto polyvinylidene difluoride membranes. Protein bands were treated with appropriate antibodies for detection and analyzed using a ChemiDoc XRS+ image analyzer (Bio-Rad). Band intensity was measured using Quantity One software (Bio-Rad). Quantitative ratios were calculated based on the data and are presented as relative values.

### Quantitative real-time PCR (qPCR) analysis

Total RNA was isolated from cells using RNAiso Plus reagent (TaKaRa), and cDNA was synthesized from 0.5 µg of each RNA preparation using the ReverTra Ace qPCR RT kit (TOYOBO) according to the manufacturer’s instructions. The following primers were used for PCR: *GAPDH*, 5′-GGA GCG AGA TCC CTC CAA AAT-3′ (sense) and 5′-GGC TGT TGT CAT ACT TCT CAT GG-3′ (antisense); and *C15ORF48*, 5′-AGG AAG GAA CTC ATT CCC TTG G-3′ (sense) and 5′-TTT TGA GGT ACA GTA GGG TCC A-3′ (antisense). After initial denaturation at 95°C for 1 min, PCR was performed for 40 cycles (15 s at 95°C and 45 s at 60°C) using a Thunderbird SYBR Green Polymerase Kit (TOYOBO) and Eco Real-Time PCR System (Illumina).

### Chromatin immunoprecipitation (ChIP) assay

ChIP assays were performed as described previously ^68^. Briefly, cells were suspended in PBS at 2 × 10^6^ cells/mL and fixed with 1% formaldehyde for 10 min at 25°C, after which fixation was halted by addition of 150 mM glycine. Fixed cells were washed with PBS and lysed with 200 mL of lysis buffer (50 mM Tris-HCl, 10 mM EDTA, and 1% SDS; pH 7.4). Cell lysates were sonicated using a bath sonicator (Bioruptor) for three cycles (power setting: high; on, 30 s, and off, 1 min; and 4°C). Sonicated lysates were centrifuged for 10 min at 10,000 × *g*, and 1,800 mL of dilution buffer (50 mM Tris-HCl, 167 mM NaCl, 1.1% Triton X-100, and 0.11% sodium deoxycholate; pH 8.0) was added to the supernatants (chromatin solutions). The chromatin solution (2 mL) was pre-cleared with 80 μL of 50% protein G-Sepharose slurry pre-absorbed with 100 μg/mL sonicated salmon sperm DNA (ssDNA), aliquoted, and incubated with 4 μL of anti-RelA antibody (Santa Cruz) or 4 μg of normal mouse IgG (MOPC21; Merck Millipore) overnight. Immunoprecipitates were recovered using 20 μL of 50% protein G-Sepharose/ssDNA for 2 h, washed sequentially with RIPA buffer (50 mM Tris-HCl, 150 mM NaCl, 1 mM EDTA, 1% Triton X-100, and 0.1% SDS; pH 8.0), RIPA buffer with 500 mM NaCl, and TE buffer (10 mM Tris-HCl and 1 mM EDTA; pH 8.0), and resuspended in 200 μL of elution buffer (10 mM Tris-HCl, 300 mM NaCl, 5 mM EDTA, and 0.5% SDS; pH 8.0). Beads and an input fraction saved before pre-clearing were incubated at 65°C for at least 6 h. DNA was extracted with phenol-chloroform, precipitated with ethanol, and suspended in 50 μL of TE buffer. Extracted DNA was subjected to qPCR analysis as described above. The following PCR primers (between −307 and −100 from the transcription start site of the C15ORF48 gene predicted from RefSeq NM_032413.4) were used:5′-GTT CCA CCT CCT ACT CCC CA-3′ (sense) and 5′-GAG GGA CCT GAC TCG CTT TC-3′ (antisense). Percent input values were calculated by comparing *Ct* values of the input and immunoprecipitated fractions and are shown as ratios relative to those of the control samples.

### Mitochondrial membrane potential determination

**M**itochondrial membrane potential was determined using tetramethylrhodamine methyl ester (TMRM) (Invitrogen). Cells were plated in 12-well plates and transfected with the indicated siRNAs for the indicated times. After incubation, cells were treated with 20 μM CCCP (Cayman Chemical) or left untreated for 6 h at 37□ and then incubated with 30 nM TMRM for 30 min at 37□. After incubation, cells were collected in 1.5 mL tubes and washed with PBS three times. The mean fluorescence intensity (MFI) of TMRM was measured using an Aria flow cytometer (BD Biosciences), and the quantitative ratio of MFI in CCCP-untreated cells to that in treated cells is shown.

### Mitophagy detection

Mitophagy was detected using a Mitophagy Detection Kit (#MD01, DOJINDO), according to the manufacturer’s instructions. Briefly, cells were seeded in a 12-well dish and incubated for 24 h. After incubation, the culture medium was discarded, and cells were washed twice with serum-free medium. Cells were then treated with mitophagy dye working solution and incubated at 37□ for 30 min. As a positive control for mitophagy, cells were additionally treated with the mitochondrial uncoupler CCCP (50 μM) for 2 h after treatment with the Mitophagy dye working solution. After incubation, supernatants were discarded, and cells were washed twice with serum-free medium and collected in a 15 mL tube. Mitophagy-dye-positive cells were detected using an Aria flow cytometer (BD Biosciences).

### ATP assay

The ATP assay was performed using an ATP assay kit-Luminescence (#A550, DOJNDO) according to the manufacturer’s instructions. Briefly, cells were plated in 96-well, white microplates and transfected with the indicated siRNAs using the reverse transfection method with Lipofectamine RNAiMax for the indicated time periods. ATP standard solutions were prepared by serial dilution with serum-free medium at 2.5, 1.25, 0.625, 0.313, 0.156, 0.078, 0.039, and 0 μM. Each diluted ATP standard solution was added to a 96-well, white microplate. After incubation, working solution was added to each well. The microplate was covered with foil, shaken for 2 min, and then incubated at 25□ for 10 min. Luminescence was measured using a Filter Max F5 microplate reader (Molecular Devices). Cellular ATP concentrations were calculated using a calibration curve.

### Glutathione assay

**G**lutathione assays were performed using a GSSG/GSH Quantification kit (#G257, DOJINDO) according to the manufacturer’s instructions. Briefly, cells were plated in a 6-cm dish and transfected with the indicated siRNAs using the reverse transfection method with Lipofectamine RNAiMax for the indicated times. After incubation, cells were collected and adjusted to 1 × 10^7^ cells/1.5 mL tube. Cell suspensions were mixed with 10 mM HCl and then lysed with two freeze-thaw cycles (30 min at -80°C and 30 min at 4°C). Five percent 5-Sulfosalicylic Acid Dihydrate (SAA) was added to samples, which were then centrifuged at 10,000 × *g* for 10 min at 4□. Supernatants were used for GSSG/GSH quantification. Standard solutions of GSSG and GSH were prepared by serial dilution with 0.5% SAA. Each diluted standard solution was then added to wells of a 96-well microplate. Buffer solution was added to each well and incubated at 37□ for 1 h. Substrate working solution and enzyme/coenzyme working solution were added to each well, and then the plate was incubated at 37□ for 10 min. Absorbance was measured at 405 nm using a Filter Max F5 microplate reader (Molecular Devices). Cellular GSSG and GSH concentrations were calculated using a calibration curve.

### MTT (3-[4,5-dimethylthiazol-2-yl]-2,5 diphenyl tetrazolium bromide) assay

Cells were plated in 96-well microplates and transfected with the indicated siRNAs using the reverse transfection method with Lipofectamine RNAiMax for the indicated time periods. After incubation, the solution in each well was replaced with DMEM containing 0.5 mg/mL MTT (DOJINDO) and incubated for an additional 1 h. The resulting crystalized product was dissolved in 100 µL of 100% dimethyl sulfoxide (DMSO) and absorbance was measured at 595 nm using a Filter Max F5 microplate reader (Molecular Devices).

### Immunocytochemistry

Images were obtained using an LSM 780 confocal laser scanning microscope with a 63×1.40 NA oil-immersion objective (Zeiss). A single planar (xy) slice (1.0-μm thickness) is shown in all experiments. Cells were fixed in 4% paraformaldehyde for 20 min at room temperature and permeabilized in PBS containing 0.2% Triton X-100 and 3% bovine serum albumin at room temperature. Cells were subsequently reacted with anti-C15orf48/NMES1 (NBP1-98391) antibody for 24 h or anti-LC3 (PM036; MBL) or anti-TOM20 (42406; Cell Signaling Technology) antibody for 1 h, washed with PBS containing 0.1% saponin, and stained with an Alexa Fluor 488- or 594-secondary antibody for 1 h. DNA was stained with a mixture of 20□µg/mL propidium iodide (PI) or 4’,6-diamidino-2-phenylindole (DAPI) and 200□µg/mL RNase A for 30□min. Stained cells were mounted with ProLong Antifade reagent (Thermo Fisher Scientific). Mitochondrial lengths and numbers of LC3 puncta were quantified using NIH image software (National Institutes of Health).

### Preparation of TEC suspensions and flow cytometry

Mouse thymi were minced with razor blades. Thymic fragments were then pipetted up and down to remove lymphocytes. Fragments were digested with RPMI 1640 medium containing Liberase (Roche, 0.05 U/mL) and DNase I (Sigma-Aldrich) and incubated at 37□ for 12 min three times. Single-cell suspensions were then stained with anti-mouse antibodies. Dead cells were excluded via 7-aminoactinomycin D (7-AAD) or SYTOX Blue staining. For staining TECs with the C15orf48 antibody (Novus Biologicals) and the analysis of Tregs, TEC suspensions and thymocytes were fixed and permeabilized with Foxp3 staining buffer set (eBioscience) according to the manufacturer’s instructions and used for intracellular staining. Cells were sorted using an Aria flow cytometer (BD Biosciences). Data were analyzed using FlowJo 10 software.

### GFP-LC3 mice analysis

For autophagy analysis of thigh skeletal muscle, *C15orf48*^−/–^/GFP-LC3 mice and GFP-LC3 control mice were maintained without food for 48 h, but with free access to drinking water. Before analysis, mice were anaesthetized and perfused through the left ventricle with 4% paraformaldehyde in PBS. Thymi were collected from normally fed mice and fixed with 4% paraformaldehyde in PBS for 4 h, followed by treatment with 15% sucrose in PBS 4 h at room temperature, and 30% sucrose in PBS overnight at 4□. Thymic samples were embedded in optimal cutting temperature (OCT) compound and stored at -80□. Sections (5-μm thickness) of the thymus in OCT compound were mounted on glass slides coated with aminosilane and fixed with ice-cold acetone for 5 min. After washing, sections were blocked with 10% goat serum and then stained with Keratin 5 or Keratin 8 together with GFP antibody (Abcam) in 10% goat serum for overnight at 4□. After washing and further incubation with fluorescence-labeled secondary antibodies for 1 h, sections were covered with glass coverslips. Images were acquired using a BZ-X710 microscope (Keyence). Areas of keratin 5-positive regions and muscle fibers and the number of LC3 puncta were calculated using BZ-X710 software (Keyence).

### Analysis of scRNA-seq

The DDBJ database of mouse thymic epithelial cells (DRA009125) was used for this analysis. Plots were created using Seurat v4.1.1 package. Cell types were defined based on expression markers as described previously ^40^.

### Immunohistochemistry

Tissues from *Rag1*^−/–^ female mice were embedded in OCT compound (Sakura) and frozen in liquid nitrogen. Cryostat sections (5μm) were fixed with ice-cold acetone for 5 min. Tissue sections were blocked with 10% goat serum and then incubated with sera from 21-week-old wild-type and *C15orf48*^−/–^ mice (100 × dilution) for 1 h at room temperature. Slides were subsequently incubated with a secondary antibody (anti-IgG-Alexa Fluor-488) and DAPI for 40 min at room temperature. Images were obtained using a confocal laser-scanning microscope (Leica).

### Quantification and statistical analysis

Graphs are presented as mean ± SD, as indicated in the figure legends. Three or more independent replicates were performed for each experiment. Student’s *t*-test was used for comparison between two groups. Comparisons of more than two groups were performed using Tukey’s multiple comparison test. Statistical significance was set at p < 0.05 (*p < 0.05, **p < 0.01, and ***p < 0.001). A p-value less than 0.05 was considered statistically significant. p-values and sample sizes can be found in the main and extended data figure legends.

