## Supplementary figures and images for "Mitochondrial protein C15ORF48 is a stress-independent inducer of autophagy that regulates oxidative stress and autoimmunity"

### Extended Data Figure 1

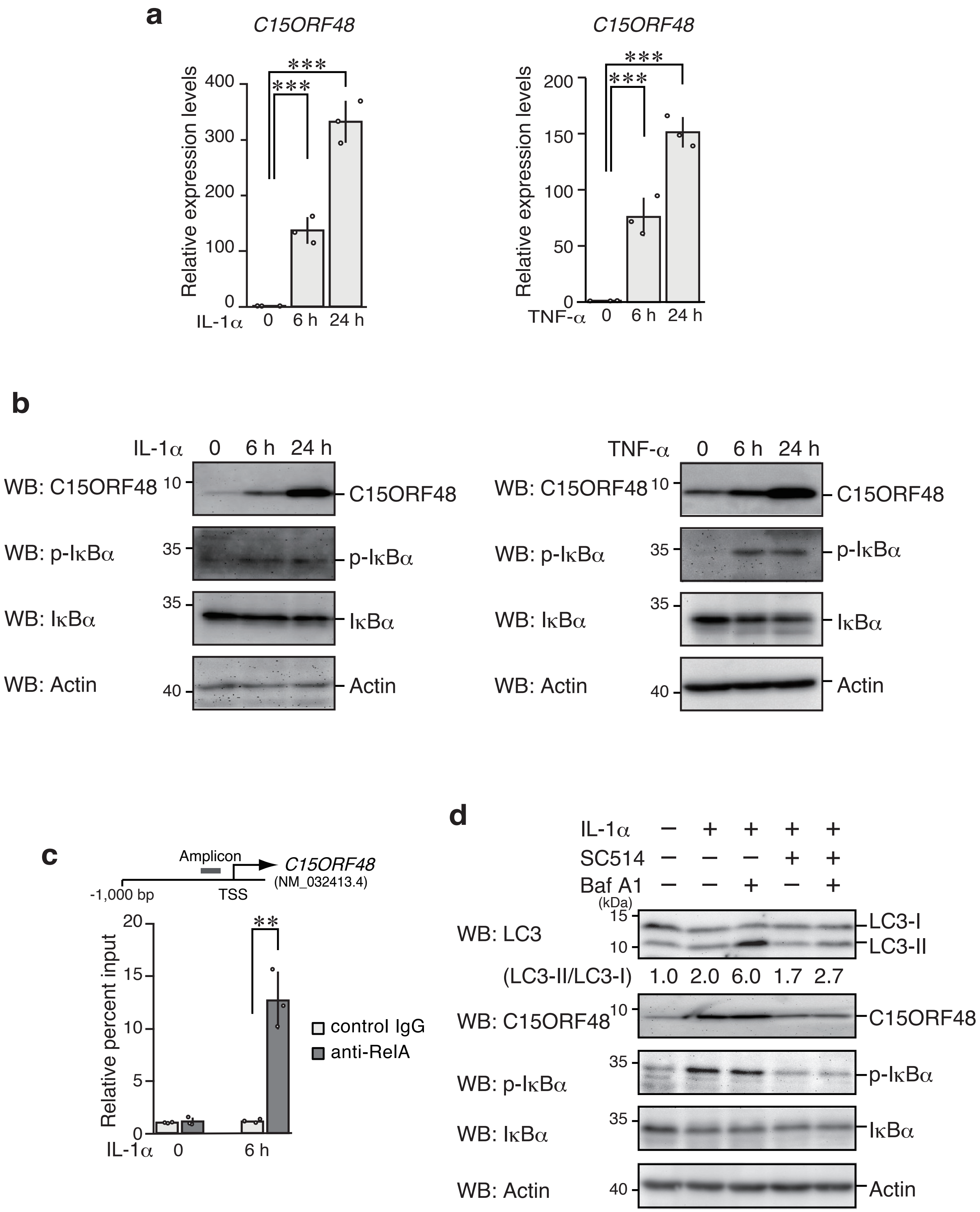


Extended Data Fig. 1

### Extended Data Figure 2

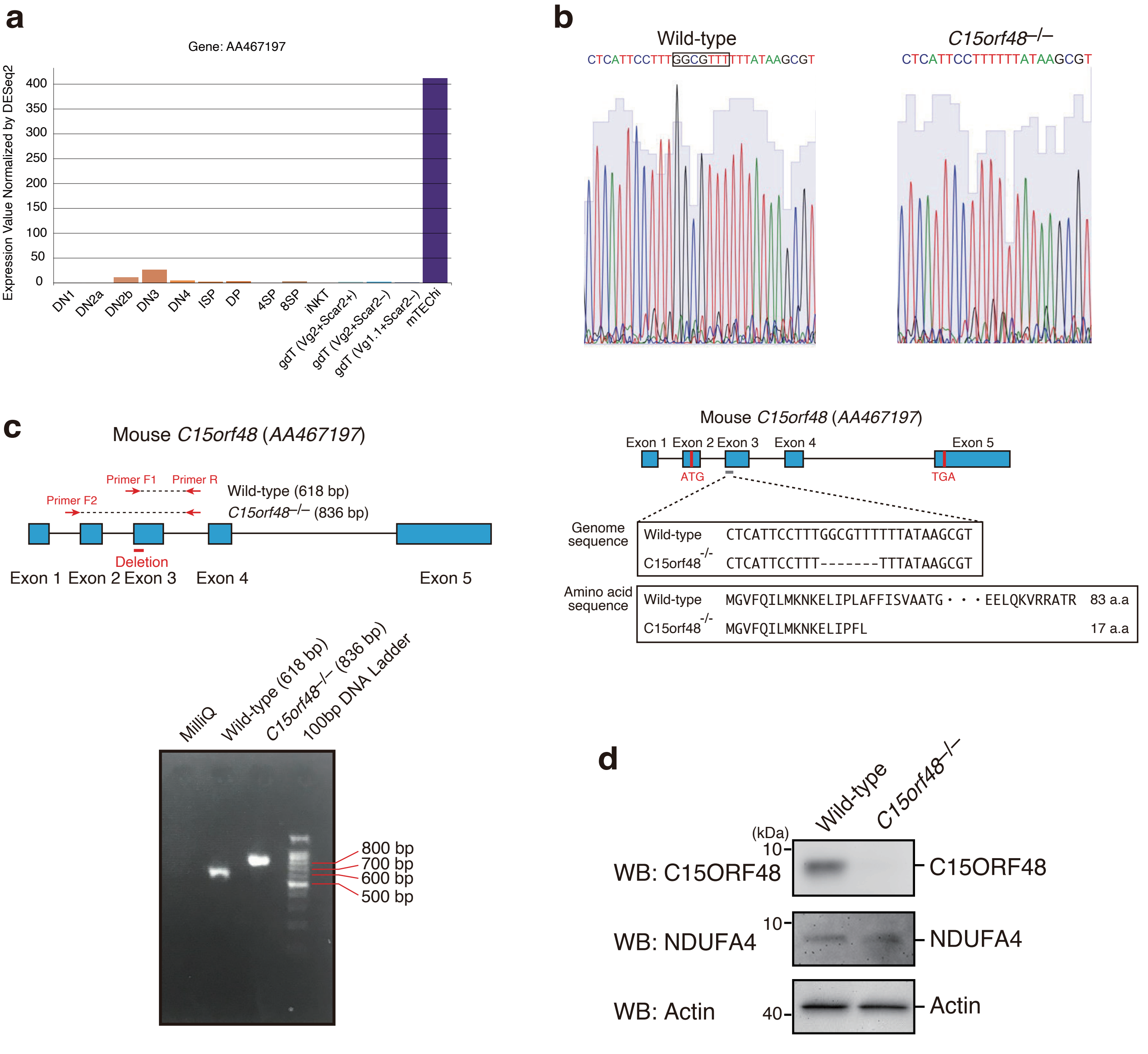


Extended Data Fig. 2

### Extended Data Figure 3

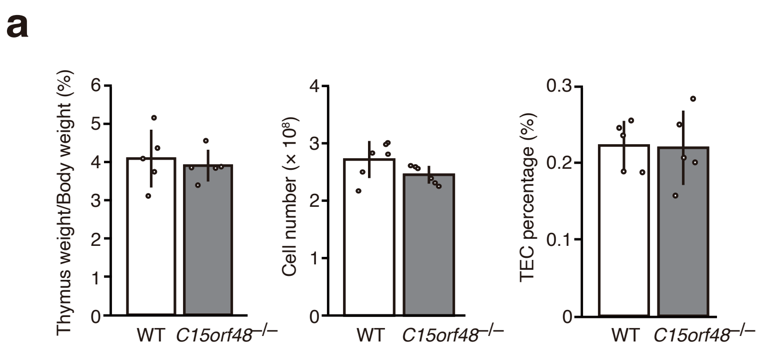


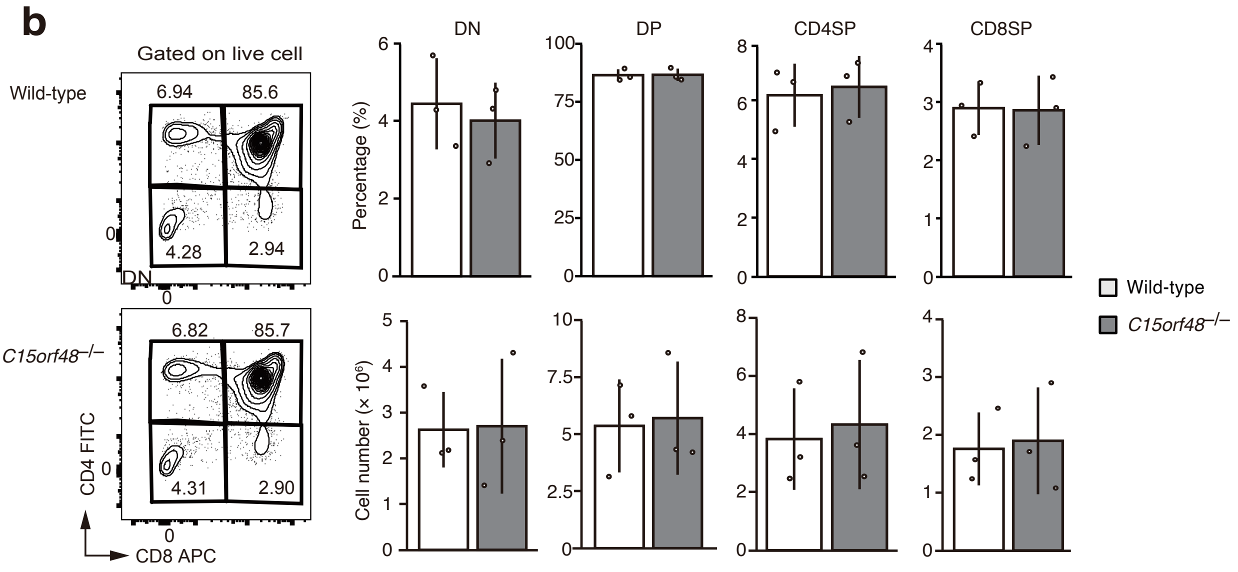


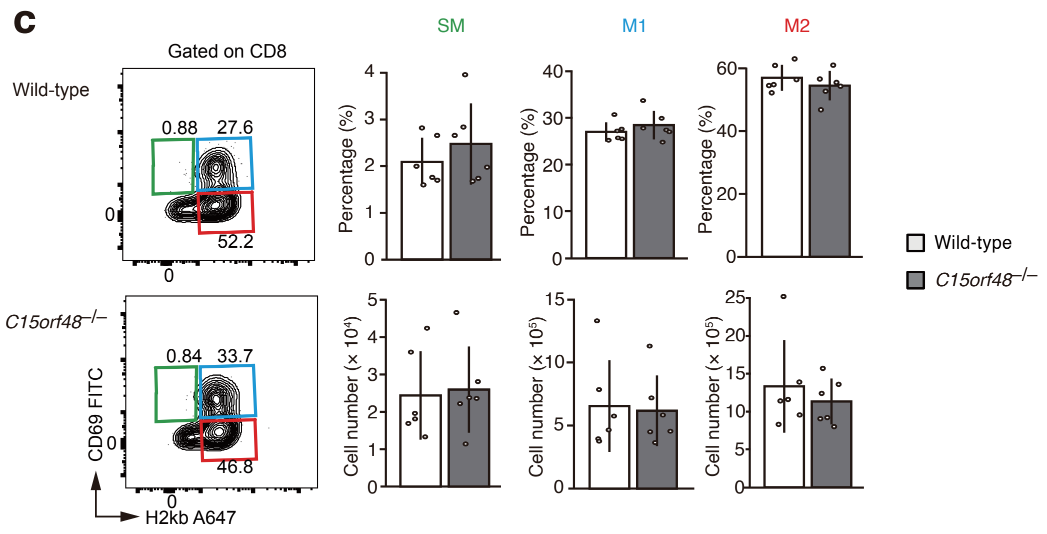


#### Extended Data Fig. 3

### Extended Data Figure 4

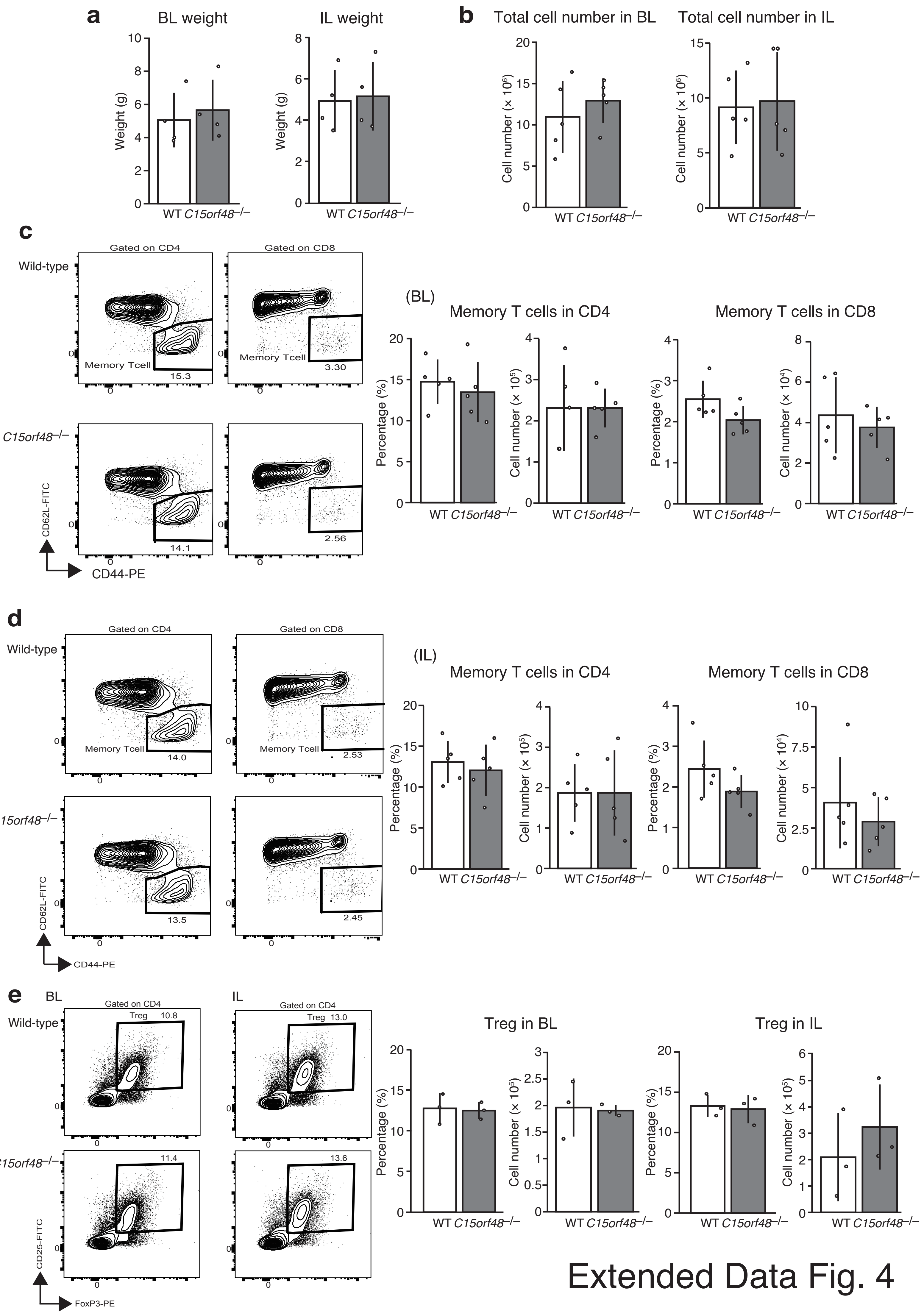


Extended Data Fig. 4

### Extended Data Figure 5

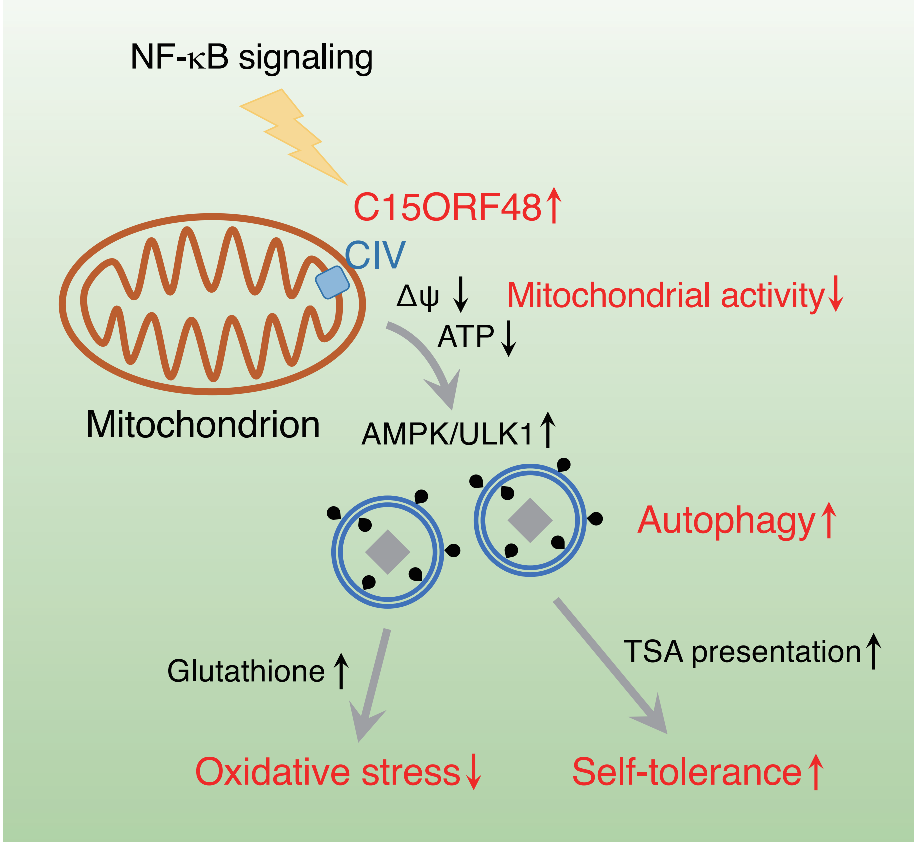


Extended Data Fig. 5
