## Supplementary Table S1 for "Mitochondrial protein C15ORF48 is a stress-independent inducer of autophagy that regulates oxidative stress and autoimmunity"

| Antibody | Source | Identifier |
| --- | --- | --- |
| Purified anti-mouse CD16/32 (Rat monoclonal) | Biolegend | Cat#101302<br>RRID:AB_312801 |
| APC/Cyanine7 anti-mouse CD45 (Rat monoclonal) | Biolegend | Cat#103116<br>RRID:AB_312981 |
| APC/Cyanine7 anti-mouse TER-119 (Rat monoclonal) | Biolegend | Cat#116223<br>RRID:AB_2137788 |
| FITC anti-mouse CD326 Ep-CAM (Rat monoclonal) | Biolegend | Cat#118208<br>RRID:AB_1134107 |
| PE/Cyanine7 anti-mouse CD326 Ep-CAM (Rat monoclonal) | Biolegend | Cat#118215<br>RRID:AB_1236477 |
| PerCP/Cyanine5.5 anti-mouse Ly51 (Rat monoclonal) | Biolegend | Cat#108315<br>RRID:AB_2632657 |
| Alexa Fluor 647 anti-mouse Ly51 (Rat monoclonal) | Biolegend | Cat#108312<br>RRID:AB_2099613 |
| PE Streptavidin | Biolegend | Cat#405203<br>RRID:N/A |
| Brilliant Violet 605 Streptavidin | Biolegend | Cat#405229<br>RRID:N/A |
| Biotinylated Ulex Europaeus Agglutinin I (UEA I) | Vector Laboratories | Cat#B-1065-2<br>RRID:N/A |
| PE anti-mouse CD80 (Armenian Hamster monoclonal) | Biolegend | Cat#104708<br>RRID:AB_313129 |
| Brilliant Violet 510 anti-mouse CD24 (Rat monoclonal) | Biolegend | Cat#101831<br>RRID:AB_2563894 |
| Brilliant Violet 785 anti-mouse Ly-6A/E (Sca-I) (Rat monoclonal) | Biolegend | Cat#108139<br>RRID:AB_2565957 |
| PE/Cynine7 anti-mouse CD4 (Rat monoclonal) | Biolegend | Cat#100528<br>RRID:AB_312729 |
| FITC Rat anti-mouse CD4 (Rat monoclonal) | BD Biosciences | Cat#553047<br>RRID:AB_394583 |
| Alexa Fluor 647 anti-mouse CD8α (Rat monoclonal) | Biolegend | Cat#100724<br>RRID:AB_389326 |
| APC/Cyanine7 a-mouse CD8α (Rat monoclonal) | Biolegend | Cat#100714<br>RRID:AB_312753 |
| FITC anti-mouse CD69 (Armenian Hamster monoclonal) | Biolegend | Cat#104506<br>RRID:AB_313109 |
| Alexa Fluor 647 anti-mouse H2-kb (Mouse monoclonal) | Biolegend | Cat#116511<br>RRID:AB_492918 |
| PE anti-mouse/human CD44 (Rat monoclonal) | Biolegend | Cat#103008<br>RRID:AB_312959 |
| PE Hamster anti-mouse γδ T-Cell Receptor (Armenian Hamster monoclonal) | BD Biosciences | Cat#553178<br>RRID:AB_394689 |
| FITC anti-mouse CD25 (Rat monoclonal) | Biolegend | Cat#102006<br>RRID:AB_312855 |
| PE anti-mouse CD25 (Rat monoclonal) | Biolegend | Cat#102007<br>RRID:AB_312856 |
| PE CD1d Tetramer | National Institute of Health | provided |
| FITC anti-mouse CD62L (Rat monoclonal) | Biolegend | Cat#104406<br>RRID:AB_313093 |
| APC anti-mouse CD357 (GITR) (Rat monoclonal) | Biolegend | Cat#126312<br>RRID:AB_2271858 |
| PE FOXP3 (Rat monoclonal) | eBioscience | Cat#12-5773-82<br>RRID:AB_465936 |
| NMES1 (C15ORF48) | Novus Biologicals | Cat#NBP1-98391<br>RRID:N/A |
| NDUFA4 | EPIGENTEK | Cat#A73444<br>RRID:N/A |
| Phospho-AMPKα (Thr172) (Rabbit monoclonal) | Cell Signaling Technology | Cat#2535<br>RRID:AB_331250 |

| | | |
|---|---|---|
| AMPKα (Rabbit monoclonal) | Cell Signaling Technology | Cat#5832<br>RRID:AB_10624867 |
| Phospho-ULK1 (Ser555) (Rabbit monoclonal) | Cell Signaling Technology | Cat#5869<br>RRID:AB_10707365 |
| ULK1 (Rabbit monoclonal) | Cell Signaling Technology | Cat#8054<br>RRID:AB_11178668 |
| Phospho-IκBα (Ser32) (Rabbit monoclonal) | Cell Signaling Technology | Cat#2859<br>RRID:AB_561111 |
| IκBα (Mouse monoclonal) | Cell Signaling Technology | Cat#4814<br>RRID:AB_390781 |
| Tom20 (Rabbit monoclonal) | Cell Signaling Technology | Cat#42406<br>RRID:AB_2687663 |
| LC3 (Mouse monoclonal) | MBL | Cat#M186-3<br>RRID:AB_10897859 |
| LC3 (Rabbit polyclonal) | MBL | Cat#PM036<br>RRID:AB_2274121 |
| ATG5 (Mouse monoclonal) | MBL | Cat#M153-3<br>RRID:AB_1278760 |
| ATG7 (Rabbit polyclonal) | MBL | Cat#PM039<br>RRID:AB_1278761 |
| NF-κB RelA (Mouse monoclonal) | Santa Cruz Biotechnology | Cat#sc-8008<br>RRID:AB_628017 |
| Actin (Mouse monoclonal) | Merck Millipore | Cat#MAB1501<br>RRID:AB_2223041 |
| α-Tubulin (Rabbit polyclonal) | Cell Signaling Technology | Cat#2144<br>RRID:AB_2210548 |
| Anti-Rabbit IgG, HRP-Linked Whole Ab Donkey | Cytiva | Cat#NA934<br>RRID:AB_772206 |
| Anti-Mouse IgG, HRP-Linked Whole Ab Sheep | Cytiva | Cat#NA931<br>RRID:AB_772210 |
| anti-GFP (Chicken polyclonal) | abcam | Cat#ab13970<br>RRID:AB_300798 |
| Purified anti-Keratin 5 (Rabbit polyclonal) | Biolegend | Cat#905504<br>RRID:AB_2616956 |
| Alexa Fluor 488 goat anti-mouse IgG(H+L) | Thermo fisher Scientific | Cat#A11029<br>RRID:AB_2534088 |
| Alexa Fluor 488 goat anti-chicken IgG(H+L) | Thermo fisher Scientific | Cat#A11039<br>RRID:AB_2534096 |
| Alexa Fluor 546 goat anti-rabbit IgG(H+L) | Thermo fisher Scientific | Cat#A11010<br>RRID:AB_2534077 |
| Normal mouse IgG MOPC21 | Merch Millipore | Cat#M5284<br>RRID:AB_1163685 |
| mTOR (Rabbit monoclonal) | Cell Signaling Technology | Cat#2983<br>RRID:AB_2105622 |
| Phospho-mTOR (Ser2481) (Rabbit polyclonal) | Cell Signaling Technology | Cat#2974<br>RRID:AB_2262884 |
| Cleaved Caspase-3 (Asp175) (Rabbit polyclonal) | Cell Signaling Technology | Cat#9661<br>RRID:AB_2341188 |
